# Specific attenuation of purinergic signaling during bortezomib-induced peripheral neuropathy

**DOI:** 10.1101/2022.02.17.479688

**Authors:** Anna-Katharina Holzer, Ilinca Suciu, Thomas Goj, Christiaan Karreman, Marcel Leist

## Abstract

Human peripheral neuropathies are poorly-understood, and the availability of experimental models limits further research. The PeriTox test uses immature dorsal root ganglia (DRG)-like neurons, derived from induced pluripotent stem cells (iPSC), to assess cell death and neurite damage. Here, we explored the suitability of matured peripheral neuron cultures for detection of sub-cytotoxic endpoints, such as altered responses of pain-related P2X receptors. A 2-step differentiation protocol, involving transient expression of ectopic neurogenin-1 (NGN1), allowed for the generation of homogeneous cultures of sensory neurons. After > 38 days-of-differentiation, they showed a robust response (Ca^2+^-signalling) to the P2X3 ligand α,β-methylene ATP. The clinical proteasome inhibitor bortezomib abolished the P2X3 signal at ≥ 5 nM, while 50-200 nM were required in the PeriTox test to identify neurite damage and cell death. A 24 h treatment with low nM concentrations of bortezomib led to moderate increases in resting cell intracellular [Ca^2+^], but signalling through transient receptor potential-V1 (TRPV1) receptors or depolarization-triggered Ca^2+^-influx remained unaffected. We interpret the specific attenuation of purinergic signalling as functional cell stress response. A reorganization of tubulin to dense structures around the cell somata confirmed a mild, non-cytotoxic stress triggered by low concentrations of bortezomib. The proteasome inhibitors carfilzomib, delanzomib, epoxomycin and MG-132 showed similar stress responses. Thus, the model presented here may be used for profiling of new proteasome inhibitors as to their side effect (neuropathy) potential, or for pharmacological studies on the attenuation of their neurotoxicity. P2X3 signalling proved useful as endpoint to assess potential neurotoxicants in peripheral neurons.

## Introduction

Models of the human peripheral nervous system are required to better understand why proteasome inhibitors (PIs) cause neuropathies. These drugs target a ubiquitous cellular function (protein degradation via the ubiquitin-proteasome system) and are used clinically to treat multiple myeloma [1, 2]. Adverse effects related to sensory neurons and nociceptors are frequent. Clinical and pathological findings include neurite damage [3–7]. However, they are also associated with several neurofunctional defects, including an altered pain regulation [8, 9]. Cell culture models for functional impariments are still very scarce.

The first proteasome inhibitor that entered clinics is bortezomib (BTZ). The boronic acid peptide reversibly blocks the chymotrypsin-like protease of the 20S proteasome [10], and it is known to induce severe adverse events in the majority of patients. Peripheral neuropathy is one of the most significant BTZ-related toxicities and affects up to 64% of patients [11–14]. BTZ-induced peripheral neuropathy (BIPN) affects long sensory neurons and the associated pain leads to therapy modification in up to 30% of the patients [12–15]. Examples for second generation PIs are delanzomib (DLZ), also belonging to the class of peptide boronic acids, and carfilzomib (CFZ), which is epoxyketone-based. Both PIs exhibit improved neurotoxic profiles, but peripheral neuropathies are still commonly experienced [16–21].

Several ion channel classes (e.g., purinergic (P2X) and transient receptor potential (TRP)) interact to regulate sensory neurons. Purinergic signaling is triggered by the binding of ATP, causing ion channels to open. Subsequent influx of cations, such as Ca^2+^ or Na^+^, leads to depolarization of the cell membrane and the generation of action potentials. In particular, the signaling via the purinoceptor P2X3 plays a role in pain perception and neuropathic pain [22–24]. P2X3, which is specifically located on the nociceptive neurons of the sensory nervous system [25, 26], contributes to the sensation of many types of pain: (i) injury-induced mechanical allodynia and thermal hyperalgesia, (ii) inflammation-induced thermal hyperalgesia, and (iii) chemical (formalin)-induced pain behaviour [23, 24, 27]. A highly complex involvement of P2X3 ion channels in pain perception is suggested by differential effects of antagonists in various pain models [23, 24].

The study of the initial mechanisms and steps leading to BIPN requires human-relevant experimental models of the peripheral nervous system. Some test methods are based on human peripheral neurons derived from induced pluripotent stem cells (iPSCs). They have been mostly used to investigate drug effects on neurite morphology or cell viability [28–31]. Such endpoints correlate with events during full-blown BIPN, such as loss of intra-epidermal nerve fibers and alterations in cytoskeletal structure and impairment of axonal transport [3–7]. Early effects of BTZ are less characterized, but they include aggresome formation (perinuclear accumulation of protein aggregates) [3] and a reorganization of the cytoskeleton in the cell somata [6]. Alterations in sensory signaling may also occur at initial stages. Despite the obvious link between neuropathic pain and abnormalities in nociceptor ion channels, only few studies focused on BTZ-induced impairments of ion channels and signaling [32–34]. It is not clear whether such findings from rodent models can be related to clinical situations, as sensory neurons of humans and other model organisms differ [35–38]. The use of human cell-based models of neuronal function may bridge this species-extrapolation gap, and provide new clues on the mechanisms underlying the initial development of peripheral neuropathies in humans.

Taking a step into this direction, we established here human iPSC-derived sensory neurons suitable for the study of altered ion channel function. We asked how well human iPSC lines differentiated towards peripheral neurons and we explored, whether transient expression of an NGN1-transgene improved the expression of functional P2X3 receptors. The usefulness of iPSC-derived sensory neuron cultures to assess PI-induced early alterations in signaling and morphology was then investigated. We focused on purinergic signaling as a sensitive endpoint affected by PIs in vitro. In parallel, microtubule arrangement in cell somata was studied as an indicator of initial morphological stress responses. Our study used a panel of five PIs to study multiple functional adaptations and to identify readouts of cell changes occuring well before signs of general cytotoxicity or a general breakdown of membrane signaling.

## Results

### Human iPSC-derived peripheral neurons for toxicity testing

Three different iPSC lines were differentiated towards peripheral neurons. The objective was to test the general applicability and robustness of a previously established two-step protocol [28, 39]. Neuronal precursors were generated and cryopreserved from the iPSC lines SBAD2, Si28 and mciPSC. After thawing and further differentiation, all cells exhibited similar neuronal morphology, neurite growth, and expression of peripheral neuron marker proteins, such as the transcription factors BRN3A and Islet-1 (ISL1), as well as the intermediate filament peripherin (PRPH). Data are displayed here for SBAD2- and Si28-derived neurons, while the process for mciPSC has been documented earlier [28] (Fig. 1A, S1). Whole transcriptome analysis of three early differentiation stages (DoD1, 4 and 7 after thawing) revealed a development of both, SBAD2- and Si28-derived neurons that was highly conserved between replicates, batches and cell lines (Fig. 1B,C). Moreover, the pattern was similar to other pluripotent stem cell lines described earlier [28]. The 50 most-regulated genes were selected and clustered: (i) the genes that were up-regulated during differentiation comprised the peripheral markers PRPH, SCN9A and RET, (ii) the genes that were down-regulated included markers for neural crest cells (PAX3, TLX2) (Fig. 1B). In a principal component analysis (PCA) of the 500 most variable genes, samples of the same differentiation stage clustered closely together irrespective of their iPSC line origin (Fig. 1C). These results confirmed that the protocol originally developed for embryonic stem cells can be broadly applied to generate peripheral neurons. In order to test, whether also functional properties were similar, we investigated toxicant-sensitivity. The PeriTox test, a well established screening assay [40–42], was used to assess effects on the neurite area and the cell viability of immature neurons on DoD0. Peripheral neurons of all three iPSC line-origins were equally sensitive to a diverse set of peripheral neurotoxicants (taxol, bortezomib, colchicine and acrylamide) (Fig. 1D). The importance of functional testing became evident, when so-called peripheral neurons were obtained from a commercial supplier. These cells reacted to colchicine and acrylamide, but they were insensitive to taxol and bortezomib (Fig. S2). Thus, these cells showed a neuronal response, as described earlier for central neurons [28]. The missing of specific peripheral toxicants (taxol, bortezomib) would make such cells unsuitable for many toxicological applications. Taken together, the two-step differentiation protocol evaluated here was found to work for a variety of iPSC lines. The high reproducibility of the peripheral neuron differentiation represents an important basis for the reliable identification of neurotoxicants in the PeriTox test. The transcriptome data obtained here provide evidence that the differentiation towards peripheral neurons continues for at least 7 days after thawing and would allow for an extension of the test period or a shift of the test window towards a more mature state.

**Figure 1:**
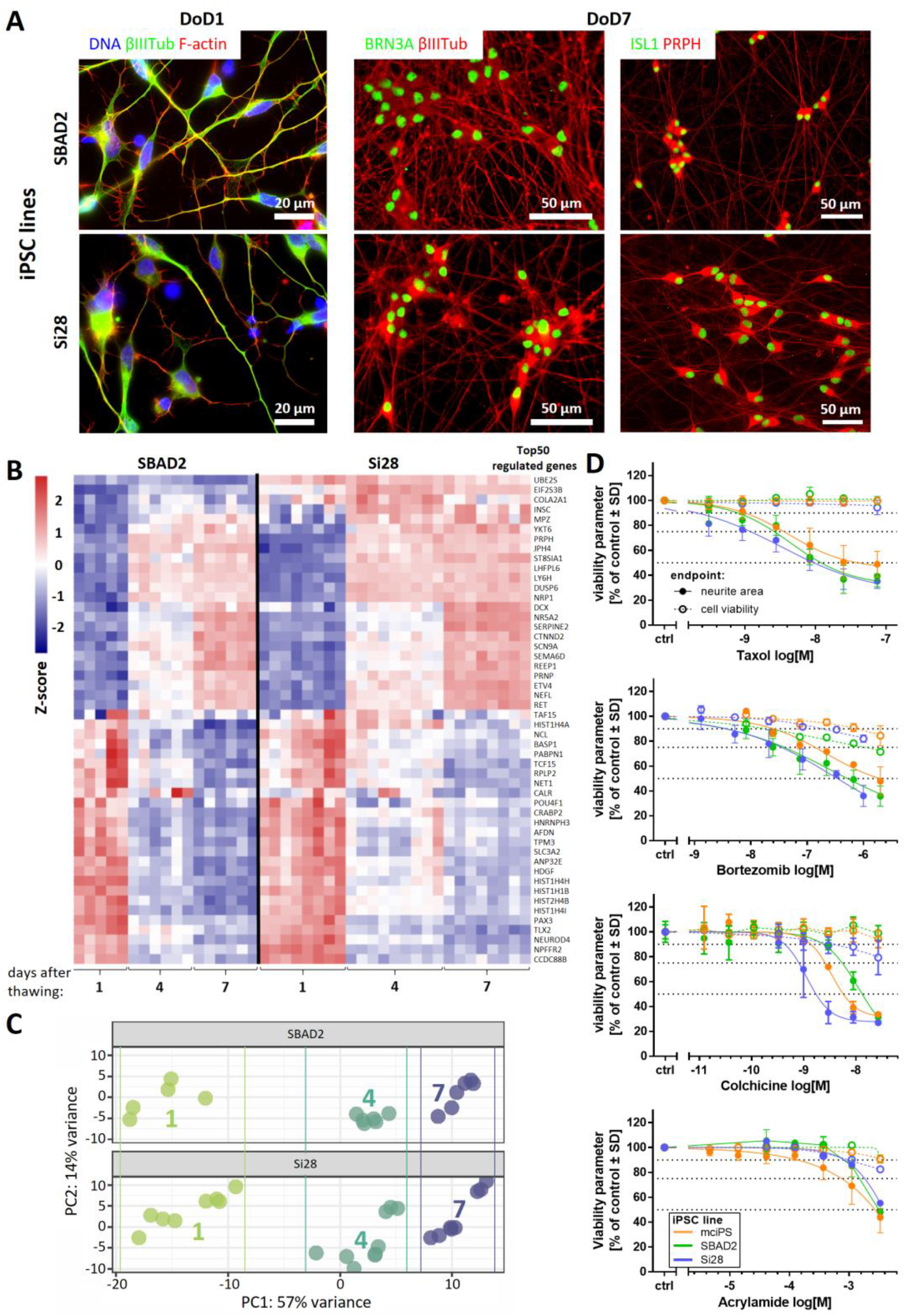
Reproducible generation of peripheral neurons from different iPSC lines and their use in the PeriTox test. **(A)** Peripheral neurons derived from the iPSC lines SBAD2 and Si28 were fixed and stained on DoD1 (left) for the neuronal cytoskeletal marker βIII-tubulin (βIIITub, green) and F-actin (red); and on DoD7 (middle, right) for the sensory neuronal transcription factors BRN3A or ISL1 (green) and the cytoskeletal proteins βIIITub or peripherin (PRPH) (red). Color code and scale bars are given in the images, details are shown in figure S1. DoDx: day of differentiation, counting from thawing of frozen neural precursors on DoD0. **(B,C)** Whole transcriptome analysis (19,000 genes) was performed for early differentiation states (DoD1, 4 and 7) of SBAD2- and Si28-derived neurons. Data are from three independent differentiations (full data in Supplementary file1). **(B)** The heatmap depicts the row-wise Z-scores of the top 50 regulated genes (exhibiting the highest variance across all samples). The upper group, defined by the clustering algorithm, mainly consists of genes up-regulated (red) during differentation and the lower group mainly consists of genes down-regulated (blue). **(C)** For the top 500 variable genes of this data set, a PCA was performed. In the two-dimensional PCA display, three differentiation stages are color-coded according to their DoD. Data points and heatmap columns correspond to all technical replicates measured in the 3 experiments per cell line. **(D)** Peripheral neurons derived from the iPSC lines mciPS (orange), SBAD2 (green) and Si28 (blue) were used in the PeriTox test. The (peripheral) neurotoxicants taxol, bortezomib, colchicine and acrylamide were used as positive controls. Effects on the neurite area (solid symbols and lines) and the cell viability (open symbols, dashed lines) are shown. Data are means ± SD of 3 biological replicates.

### Need for novel test strategies to further improve sensitivity

Although the PeriTox test has been used successfully to screen for environmental chemicals, an increased sensitivity is desirable for pre-clinical testing of drugs. To refine the standard PeriTox test, scenarios of prolonged exposure to toxicants at different time points of differentiation were investigated (Fig. 2A, S3A). First, a prolonged toxicant exposure time (48 h and 72 h, DoD0-2 and DoD0-3, respectively) was explored. The sensitivity of neurites to acrylamide, colchicine and taxol did not change significantly. However, the rate of cell death increased with prolonged incubation time (Fig. S3B-D). For bortezomib, the neurite area was affected more with longer exposure times. However, this effect was attributable to the concomitant decrease in cell viability (Fig. 2B, S3E). Taken together, these findings meant, that the assay became less specific for neurite toxicants. The prediction model for the standard PeriTox test [28, 42] requires the specific toxicants to affect neurites at concentrations three times lower than cell viability. This requirement was not met in the prolonged assay (Fig. 2C,D).

**Figure 2:**
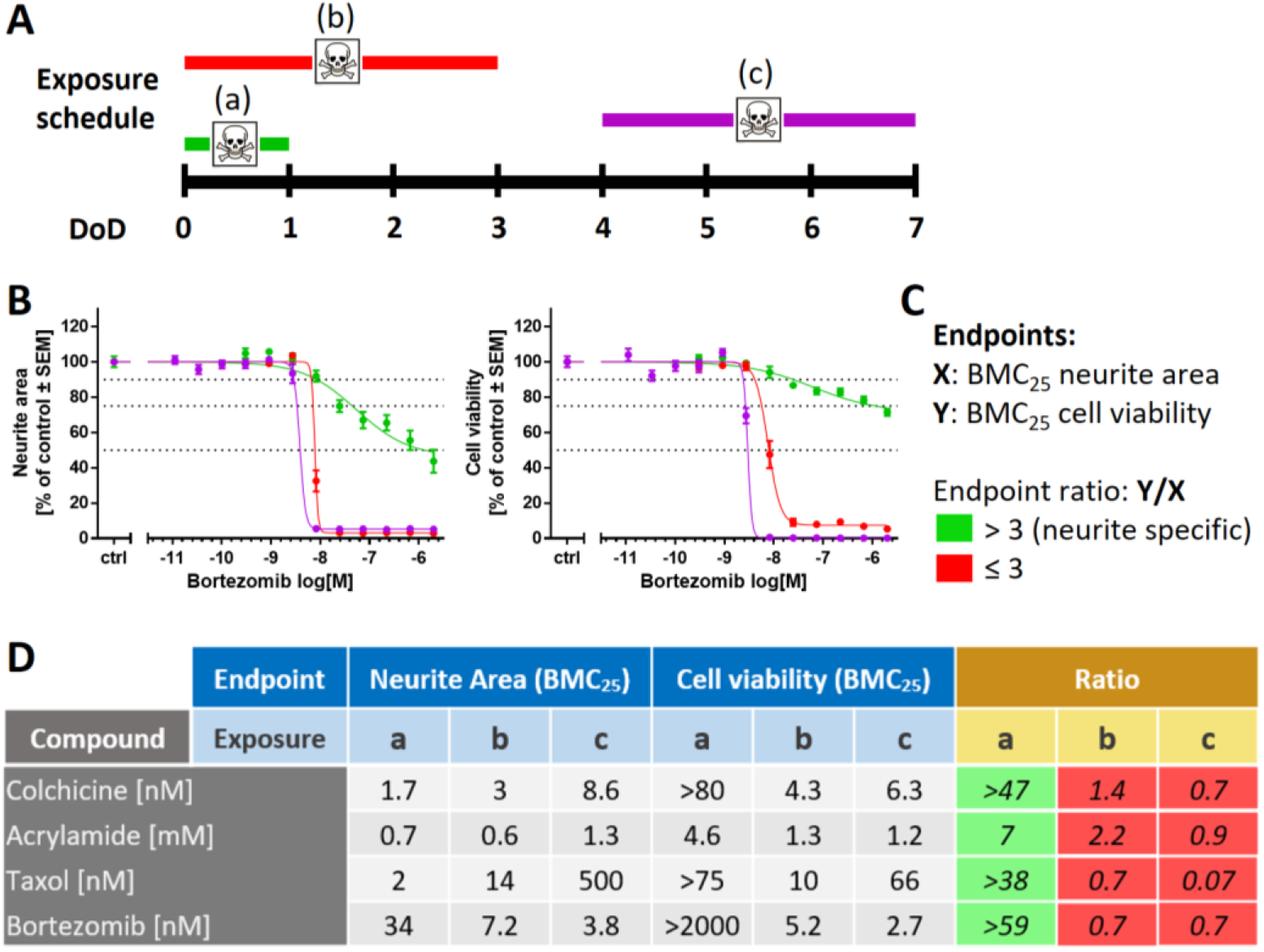
Variation of the exposure schedule to assess compound toxicity. **(A)** Schematic representation of the applied exposure schedules with a 24 h treatment starting on DoD0 (a, standard PeriTox test, green), immediate 72 h treatment (b, DoD0-3, red) and delayed 72 h treatment (c, DoD4-7, purple). DoDx: day of differentiation, counting from thawing of frozen neural precursors on DoD0. **(B)** SBAD2-derived peripheral neurons were exposed to bortezomib according to the three exposure schedules. Effects on the neurite area and the cell viability were assessed. Data are means ± SEM of 3 independent experiments. **(C)** Prediction model for the classification of compound-induced effects: The concentrations relating to the benchmark response level of 25% decrease of a test endpoint (BMC25) was calculated for both endpoints, neurite area (X) and cell viability (Y). A ratio of Y/X > 3 is classified as a “neurite-specific” compound effect (green); Y/X≤ 3 marks effects that are “not neurite-specific”, such effects were classified as “cytotoxic” (red). **(D)** BMC25 values were calculated for both test endpoints in all three exposure scenarios. Effects induced by colchicine, acrylamide, taxol and bortezomib were classified according to the prediction model. Respective concentration-response curves are given in figure S4.

Next, we explored, whether shifting the time window of exposure to a later time point (DoD4-7) would result in more potent neurite toxicity. This was not the case (Fig. 2B,D, S4A,B). Moreover, any specificity for neurite effects (relative to general cell death) was lost. Altogether, these results suggest that polonged toxicant exposure is not a suitable measure to improve the PeriTox test. We concluded that other approaches and new functional endpoints are required for a more sensitive assay for compounds that may trigger peripheral neuropathies.

### Purinergic signaling as a functional feature of iPSC-derived sensory neurons

One of the most important functional changes during peripheral neuropathy is altered pain perception. This suggests that assessment of pain-related neuronal signals might be a suitable endpoint for peripheral neurotoxicity testing *in vitro*. To explore this possibility, we set out to generate cultures of peripheral neurons that allowed the quantification of nociceptor function.

Our preliminary experiments showed that peripheral neurons could be cultured and further matured for at least 2 months. However, we did not succeed in obtaining robust nociceptor responses suitable for drug screening. For this reason, we introduced an inducible NGN1 transgene into the iPSC line Si28 to generate Si28-NGN1 cells. This strategy has been described earlier to enhance nociceptor differentiation [43], and we found indeed that our 2-step protocol, enhanced by induction of NGN1 for a defined time period, led to an improved differentiation. The neurons generated by this protocol (Fig. 3A) were found to be post-mitotic already on DoD1 after thawing (Fig. 3B, S5A,B). They could be cultured for at least 42 days as stable neuronal network suitable for single cell observations (Fig. 3C, S5C). To characterize the Si28-NGN1-derived neurons, we tested them for the expression of the nociceptor-specific receptors P2X3 and TRPV1. Immunostaining showed that most cells (>80%) were P2X3 and peripherin double-positive (Fig. 3D, S5D). Moreover, we used differentiated neurons in Ca^2+^-imaging experiments: Cells were generated from the iPSC lines SBAD2 and Si28 as well as Si28-NGN1 and used after at least 38 days of differentiation. Only <15% of neurons fom standard iPSCs responded to the P2X3-specific agonist α,β-methylene ATP (α,β-meATP). More than 80% of the neurons generated from Si28-NGN1 revealed increased [Ca^2+^]_i_ upon application of an α,β-meATP stimulus (Fig. 3E). The transient signal in continued presence of the ligand was typical for self-inactivating P2X3 ion channels (Fig. 3F). The application of the TRPV1-specific agonist capsaicin hardly stimulated neurons from SBAD2 or Si28. About 40% of all neurons in cultures from Si28-NGN1 showed a clear response. This was specifically blocked by a TRPV1 antagonist (Fig. S6). The sub-population responding to ATP (a general agonist for all P2X receptors) was of similar size to the P2X3-responsive neuronal sub-population. Moreover, the strong efficacy of a P2X3-specific antagonist to block ATP responses suggested that most functional P2X receptors were P2X3 (Fig. 3E,F).

**Figure 3:**
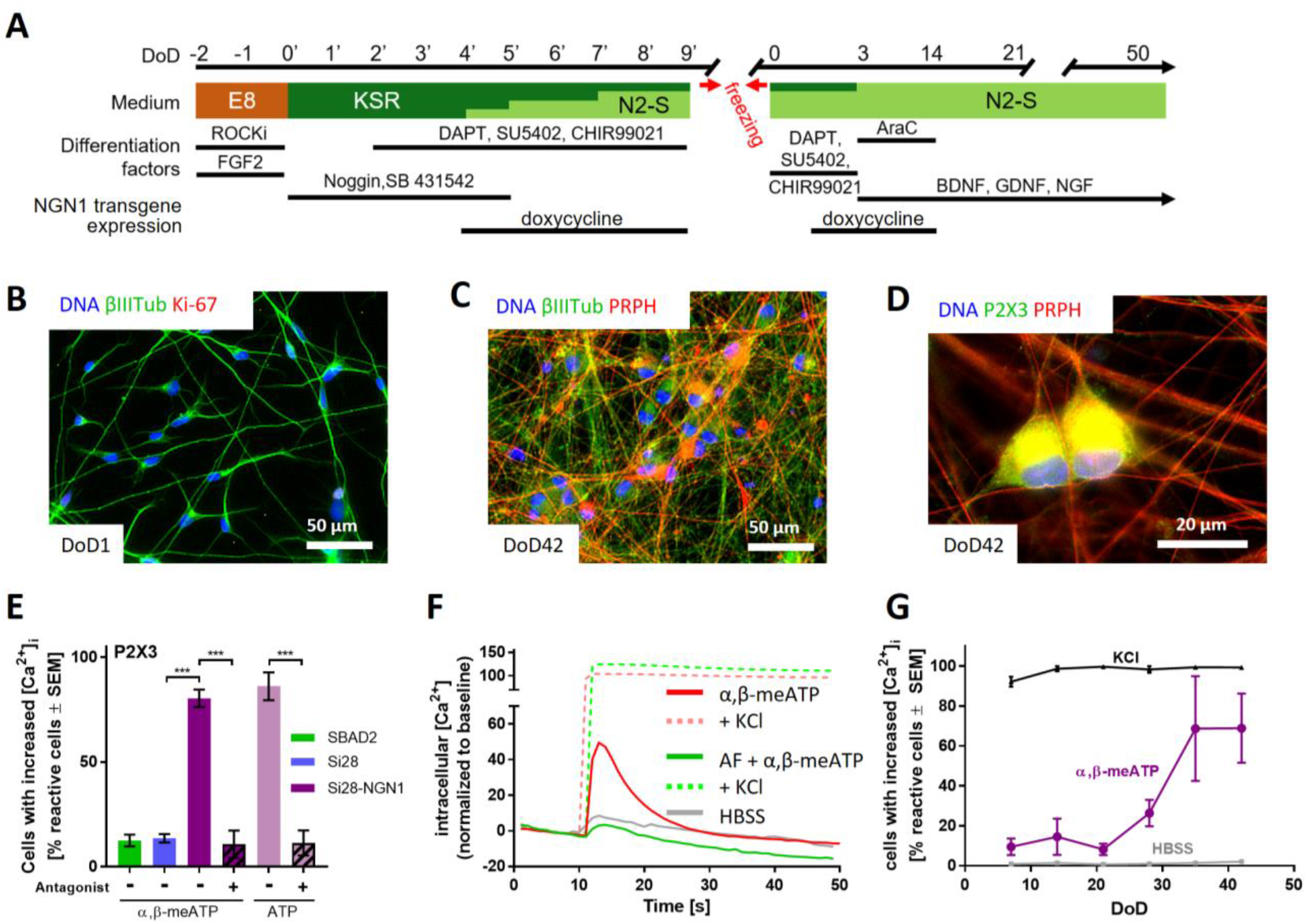
Sensory neurons exhibiting functional P2X3 receptor signaling. **(A)** Schematic representation of the differentiation protocol for the generation of functional sensory neurons from the genetically modified iPSC line Si28-NGN1. During the standard differentiation procedure, transient NGN1-transgene expression was induced from DoD4’ until DoD9’ and from DoD1 until DoD14 by addition of doxycycline. DoDx’: day of differentiation, counting from pluripotent state (DoD0’); DoDx: day of differentiation, counting from thawing of frozen neural precursors on DoD0. Other factors added (e.g., ROCKi) are detailed in the methods. **(B-D)** Representative immunofluorescence images of cells fixed on DoD1 and stained for αIII-tubulin (αIIITub) and the proliferation marker Ki-67 **(B)**; or on DoD42 and stained for peripherin (PRPH) and αIIITub **(C)** or P2X3 **(D)**. Nuclei were stained using H33342 (DNA). Color code and scale bars are given in the images. Details are shown in figure S5. **(E)** Peripheral neurons derived from the iPSC lines SBAD2 (green), Si28 (blue) and Si28-NGN1 (purple) were differentiated for >38 days and used for Ca^2+^-imaging experiments. The P2X3-specific agonist α,ß-methylene ATP (α,β-meATP) was used to determine the expression of functional P2X3 receptors. ATP was used as a general agonist for purinergic receptors. AF-353, a P2X3-specfic antagonist, was used to confirm exclusive P2X3 expression. **(F)** Exemplary traces (red) of changes in intracellular [Ca^2+^] upon α,β-meATP (1 μM) application (solid lines). After the primary stimulus, KCl (dashed lines) was added. Some cells were pre-treated with AF-353 (0.1 μM) (green). The grey line depicts changes upon application of the negative control (HBSS, grey). **(G)** Time-dependency of the expression of functional P2X3 receptors. Sensory neurons were tested weekly for their potential to respond to HBSS, α,β-meATP and general membrane depolarization induced by KCl. **(E,G)** Data are means ± SEM of 3 independent biological replicates. ***p < 0.0001.

For experimental logistics, it is important to know how long cells need to be differentiated to reach good functionality. Therefore, neurons were tested after increasing differentiation times: at DoD7, already >90% of neurons showed a Ca^2+^-response upon depolarization (KCl), but no response to P2X3 stimulation. The latter response started to increase at DoD20-30, and reached its saturation level at >DoD35 (Fig. 3F).

Taken together, the overexpression of NGN1 during early differentiation steps allowed the generation of <u>p</u>eripheral <u>n</u>eurons with enhanced <u>n</u>ociceptor features (<u>PNN</u>). PNN were found suitable to quantitatively evaluate functional P2X3 responses of single cells in Ca^2+^-imaging experiments. Next, a full transcriptomic characterization of this promising drug discovery model was performed.

### Transcriptomics profile of Si28-NGN1-derived sensory neurons

The expression levels of about 19,000 genes were determined for 6 differentiation stages of sensory neurons generated from Si28-NGN1 cells (Suppl. File2). A principle component analysis (PCA) of the whole set of genes provided a first overview on the dynamics of gene expression, and showed a continuous progression of cell differentiation until DoD42. Furthermore, the PCA demonstrated the good reproducibility of the differentiation protocol, as three independent differentiations clustered closely together (Fig. S7A). Transcriptome changes continued until late differentiation stages (DoD35-42) as shown by the up-regulated gene expression of, e.g., plexin C1 (*PLXNC1*), which is involved in axon guidance, and the serotonin receptor 2A (*HTR2A*) [44, 45], and the down-regulation of growth cone-related genes, such as *ROBO2*, and the netrin receptor *UNC5B* [46, 47] (Fig. S7B).

To generate a condensed overview of the expression profile for Si28-NGN1-derived PNN, a small panel of 122 genes characteristic for neural cell types and signaling pathways was assembled (Fig. 4). Most pan-neuronal markers included in this panel were found to be expressed already on DoD1 and remained highly expressed over 6 weeks (e.g., neurofilaments (*NEFL/M/H*), acetylcholine esterase (*ACHE*) and microtubule associated protein tau (*MAPT*)). The sensory neuronal marker genes *ISL1, POU4F1* (BRN3A) and *PRPH*, the nociceptor markers *SCN9A* and *RET*, as well as various pre- and post-synaptic markers showed high expression levels throughout the monitored time of differentiation. Neural crest-specific genes (i.e. those related to PNN precursors), such as *PAX3* and *MSX1*, were down-regulated over time. These data confirm that the newly established differentiation protocol yields peripheral neurons with many features expected from nociceptors. Relatively few indications for other cell types were found, as only a subset of potentially glial genes was expressed, and there was little evidence for non-neural cell types.

**Figure 4:**
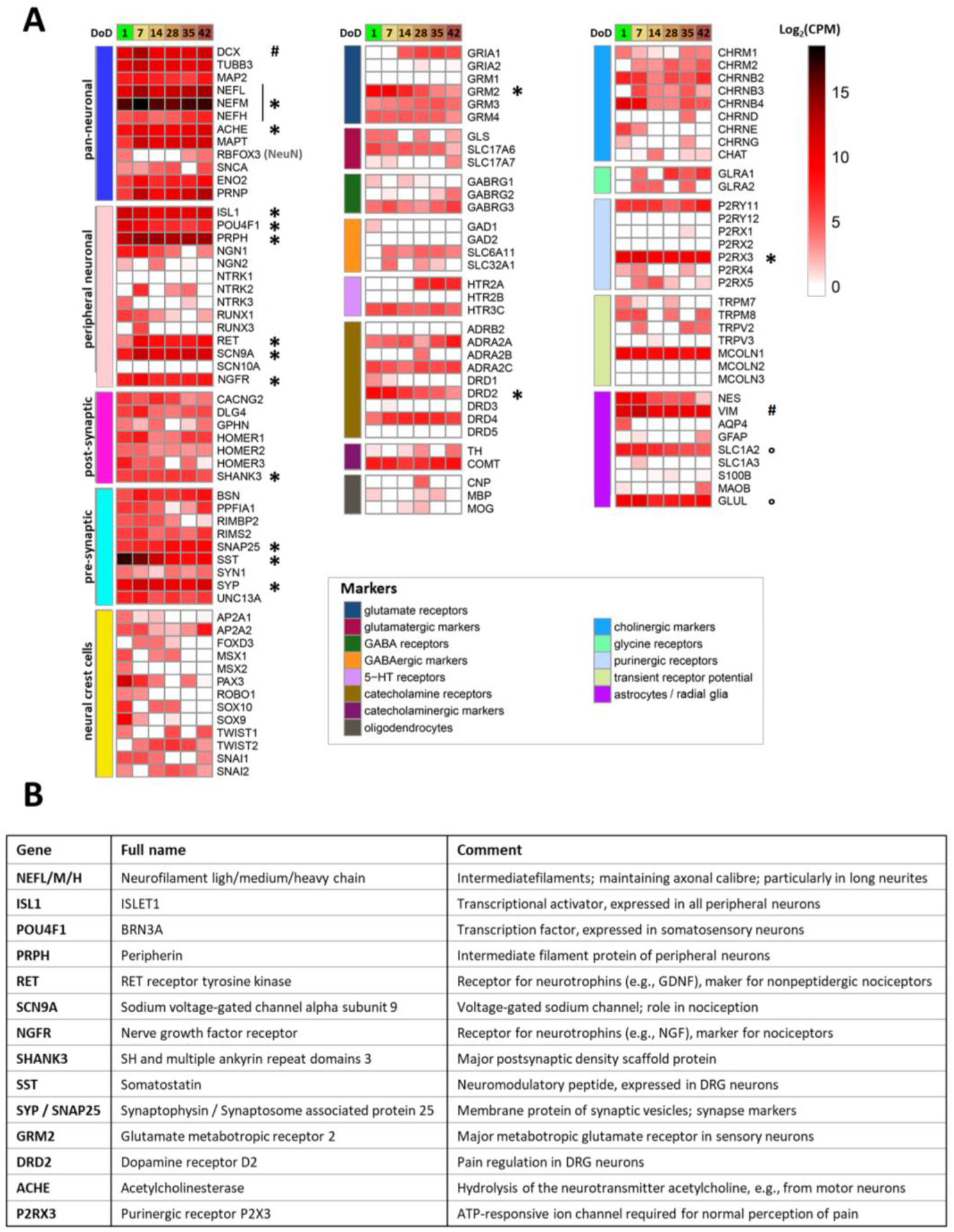
Transcriptome profiling of Si28-NGN1-derived sensory neurons. Neurons were pre-differentiated to immature sensory neurons and frozen. **(A)** After thawing, gene expression levels were determined for 6 differentiation stages (on day of differentiation (DoD) 1, 7, 14, 28, 35 and 42) by the TempO-Seq method. The heatmap visualizes the normalized counts for each gene (rows) and the DoD (columns). The neuronal overview panel of 122 genes is clustered by gene groups (e.g. neuronal and glial subtypes, receptor and ion channel classes). The gene groups are indicated by color bars (left). The absolute expression levels are given in counts of the corresponding gene per 1 million reads (CPM). The color scale uses log2(CPM) units (see supplementary files for complete data sets) and ranges from white (no expression) to dark red (high expression). Data are derived from 3 independent differentiations. A subset of genes that that may be used for routine culture controls is highlighted (*). High expression levels of VIM and DCX (#) indicate a still relatively “young” state of the cells that may be even further matured. SLC1A2 and GLUL (°) are often considered glial markers, but the absence of GFAP, AQP, S100B and MBP indicate that the cultures do not contain classical astrocytes or Schwann cells. **(B)** Overview of highly expressed differentiation markers highlighted in **(A) (*)**, with their full names and a brief explanation of their biological functions.

Especially the pattern of receptor subtypes was highly distinct, as indicated here by three examples.: (i) Amongst dopamine receptors,the D2 subtype (*DRD2, DRD4*), which is known to be expressed in dorsal root ganglia (DRG) neurons [48] was dominant, whereas *DRD3* and *DRD5* transcripts were absent; (ii) Genes encoding the metabotropic glutamate receptors 2 and 3 (*GRM2/3*), both expressed in human DRG neurons [49, 50], were found to be expressed, but not *GRM1*; (iii) Among the P2X receptors, only the nociceptor-characteristic P2X3 transcripts were measured at all differentiation stages.

In a last step, we picked a limited set (n=17) of highly expressed genes (Fig. 4A). We felt that these genes could be suitable for differentiation control by PCR for further use of the cultures or for inter-laboratory method transfer. A brief overview of the broad biological functions covered was assembled (Fig. 4B). In this context, it was interesting to see that *RBFOX3* mRNA levels were relatively low. This gene codes for the pan-neuronal marker NeuN that is very frequently used for immunostaining of CNS neurons by the community [51–53]. The low gene expression in PNN was consistent with our finding that these cells very poorly stain for NeuN (not shown), compared to all our other central neuronal cultures [39, 54, 55].

The transcriptome analysis confirmed that even after more than 30 days of PNN cultivation, the differentiation processes are not fully completed. Ongoing alterations at the level of gene expression may explain why P2X3 responses of PNN are observed only at ≥DoD28 (Fig. 3G). For documentation of late transcriptome changes, we compiled exemplary genes that are clearly (> 4-fold) and significantly regulated at late time points (DoD35-42) relative to DoD7 (Fig. S7B). These observations supported our decision to use PNN for further functional studies at late stages of differentiation, i.e. at >DoD35, to ensure the best possible maturation.

### Purinergic signaling as test endpoint to assess peripheral neurotoxicity

To explore the usefulness of Ca^2+^-imaging as a readout for disturbed pain signaling, we first investigated two clinically used proteasome inhibitors (PIs) known to cause peripheral neuropathy: bortezomib and carfilzomib. Pre-screening of the compounds in the PeriTox test indicated a cytotoxicity threshold of 200 nM for bortezomib and 66 nM for carfilzomib (Fig. 5A, left, middle). PNN were exposed on DoD≥38 to sub-cytotoxic concentrations (5 and 20 nM) for 24 h. After this “drug treatment”, we tested whether the neurons were still able to show purinergic signaling. Bortezomib concentrations of 5 nM and higher resulted in a complete shut-down of P2X3 signaling, as indicated by Ca^2+^-imaging experiments (Fig. 5B, left, S8B). Carfilzomib induced a similar non-responsiveness at ≥20 nM (Fig. 5B, middle). In order to make sure that neurons were not made generally non-responsive by a cytotoxic response missed in the PeriTox test, they were exposed to a membrane depolarizing KCl stimulus after the α,β-meATP stimulation. The cells still showed Ca^2+^-flux at PI drug concentrations that had blunted P2X3 signaling (Fig. S9A). Thus, neurons were still able to respond by Ca^2+^-signaling, and we suggest that PI treatment specifically impairs purinergic signaling. As further control, we investigated signaling through pain-related TRPV1 receptors. PI-treated neurons did not differ from control cells in this response (Fig. S9B). These results further confirm that the attenuation of P2X3 signaling was not attributable to generally decreased cell viability, or an overall loss of signaling functions.

**Figure 5:**
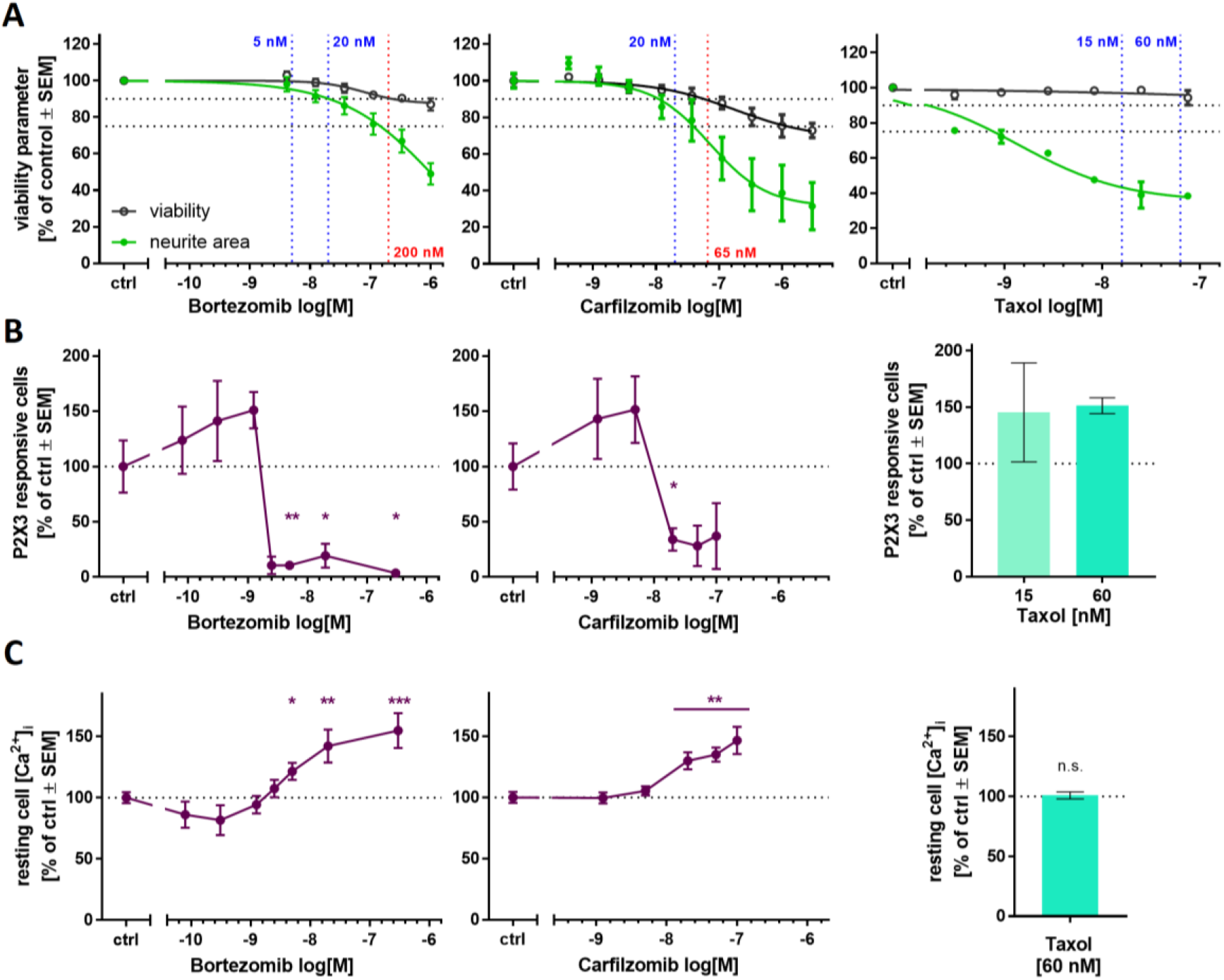
Ca^2+^ signaling as sensitive functional endpoint to assess proteasome inhibitor toxicity. The compounds bortezomib (left), carfilzomib (middle) and taxol (right) were investigated regarding their effects on different test endpoints. **(A)** The PeriTox test was used to assess their effects on neurite area and viability. Horizontal dashed lines at 90% and 75% indicate the cytotoxicity threshold the neurite effect threshold, respectively. Vertical dashed lines indicate the lowest cytotoxicity-inducing concentration (red) and the concentrations further used for Ca^2+^-imaging experiments (blue). **(B,C)** Sensory neurons (>DoD38) were pre-treated with the test compounds for 24 h, before Ca^2+^-imaging experiments were performed. **(B)** The number of cells responsive towards stimulation with the P2X3-specific agonist a,β-methylene ATP was assessed. **(C)** Baseline fluorescence, indicating the resting cell intracellular [Ca^2+^]_i_ was quantified for whole sensory neuron cultures. Exemplary single cell fluorescence traces are shown in figure S9. **(A-C)** Data are given as % of untreated control cells and are means ± SEM of at least 3 biological replicates. * p < 0.05, ** p < 0.001, *** p < 0.0001.

On closer inspection, we observed that pre-treatment with bortezomib or carfilzomib led to a mild deregulation of [Ca^2+^]_i_ in the unstimulated state (Fig. 5C, S8A). This may explain an unresponsiveness of P2X3, possibly as counter-regulation or tachyphylaxis mechanism.

To address the question of whether also a non-PI peripheral neurotoxicant would attenuate P2X3 signaling, we repeated several of the above experiments with taxol. The chemotherapeutic drug group of taxanes (including taxol) alters microtubule dynamics, but does not affect the proteasome function. Exposure to taxol in the PeriTox test showed no effect on cell viability at concentrations up to 75 nM, but neurites were strongly affected at concentrations ≥1 nM (Fig. 5A, right). We chose pre-treatment conditions of 15 and 60 nM to test for functional impairments of P2X3 or TRPV1 receptors and of depolarization induced Ca^2+^-influx. None of the endpoints was affected (Fig. 5B, right, Fig. S9). These findings suggest that impaired P2X3 signaling is a sensitive and specific endpoint for early PI-induced functional impairments.

### PI-associated reorganization of the microtubule structure in cell somata

We used several structural endpoints to potentially identify additional features of mild cell stress that would parallel impaired P2X3 signaling in the low nM range. We hypothesized that such findings would give additional evidence for early non-cytotoxic changes that preceede full-blown neuropathies. Staining of PNN for the cytoskeletal protein βIII-tubulin confirmed that the neurite network was fully intact (no neurite fragmentation or blebbing). However, we observed a conspicuous ring-like tubublin accumulation in the periphery of cell somata of bortezomib- and carfilzomib treated cells (Fig. 6A, S10). To follow up on this, cells exhibiting such a circular tubulin structure were quantified. Distinctive microtubule reorganization occurred in >80% of the cells pre-treated with PI concentrations that also resulted in the attenuation of P2X3 signaling (Fig. 6B). Further experiments showed that the accumulation in ring structures was a tubulin-specific phenomenon, as such structures were not found in stainings of the same cells for the cytoskeletal intermediate filament peripherin (Fig. S10). However, also peripherin showed a mild reorganization phenotype: While its structure in neurites was not altered, PI-treated cells showed some peripherin clustering in the somata. This was mainly seen in cytosolic areas (outside the nucleus, but not ring-shaped under the plasma membrane). For comparison, PNN were also treated with taxol (60 nM). A small number of cells presented with tubulin accumulations (Fig. 6B). Closer examination revealed that these structures were more diffuse than the very sharp rings triggered by PIs (Fig. 6A, S10). Thus, sharp tubulin-rings correlated with P2X3 impairment. These findings are in good agreement with observations in primary dorsal root ganglia that accumulation of cytoskeletal proteins in the cell somata is specific for early PI-induced neuronal stress [6, 56].

**Figure 6:**
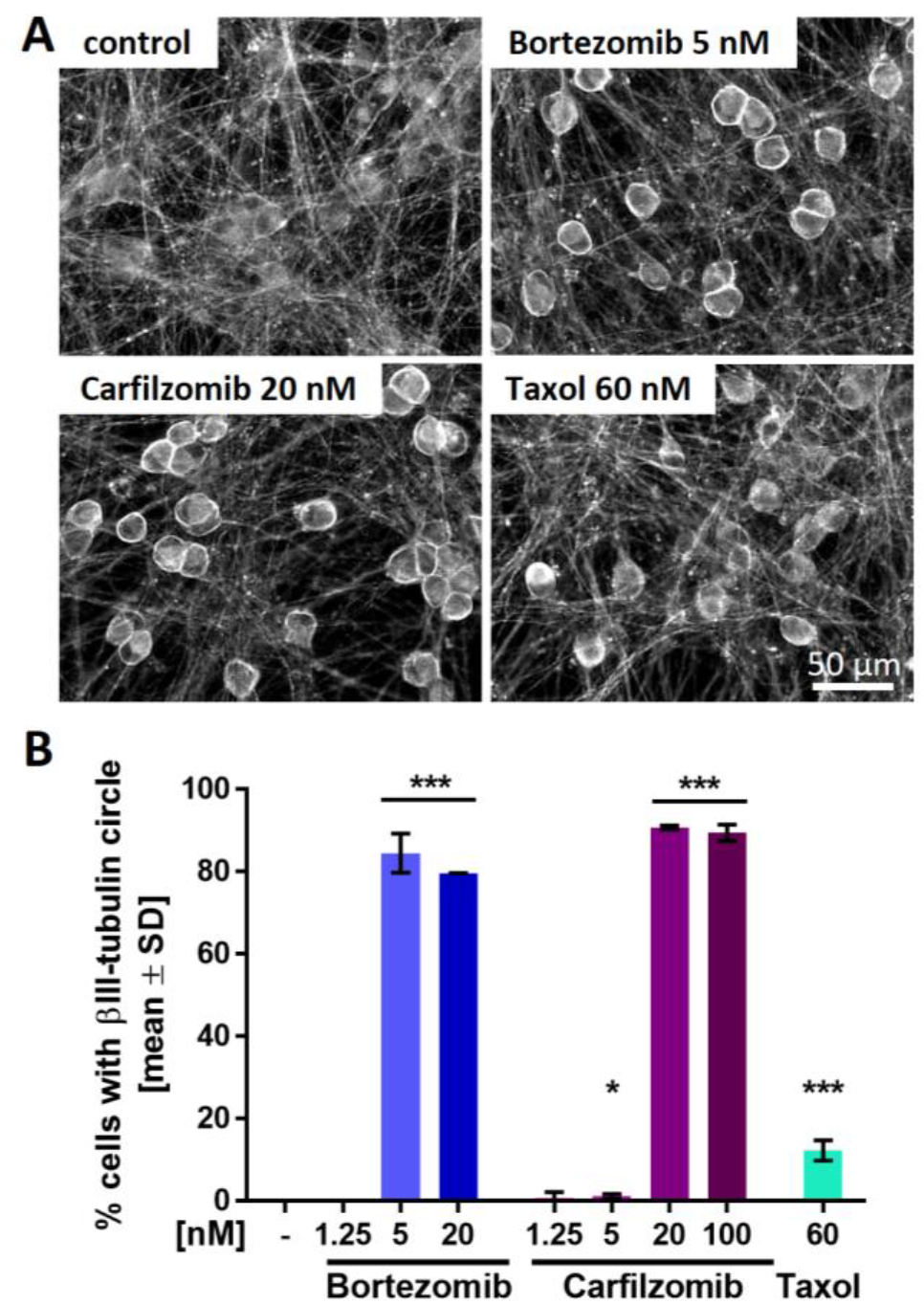
Proteasome inhibitor-induced reorganization of the microtubule structure in neuronal somata. Sensory neurons were differentiated for at least 38 days after thawing and exposed to bortezomib, carfilzomib or taxol for 24 h before fixation. (A) Representative immunofluorescence images of cells stained for αIII-tubulin. Scale bar is given in the images, and further details are shown in figure S11. (B) Cells exhibiting intense, circular αIII-tubulin staining around the cell somata (covering at least 50% of a full circle) were quantified. Data are given as % of the total cell count (number of viable cell nuclei) and are means ± SD of 2-3 biological replicates. * p < 0.05, *** p < 0.0001

While we studied the accumulation of cytoskeletal elements in somata, we wondered whether PNN nuclei were also affected by PIs. The neurons were examined in more detail for signs of condensed or fragmented chromatin, indicative for apoptotic cells. No changes in the size of neuronal nuclei or the intensity of the DNA stain were observed. However, the nuclei had an altered (more bean-shaped) morphology (Fig. S11). This may be a consequence of protein accumulations in the cytosol exerting “pressure” on the normally more rounded nuclei.

Taken together, these data show that P2X3 impairment was accompanied by a structural change, i.e., cytoskeletal protein accumulation in somata. This occurred at concentrations that did not alter any other endpoint investigated in this study. In the next step, we investigated whether our findings applied to PIs in general.

### Blunted P2X3 signaling and tubulin re-organization as PI class-effects

To explore whether impairment of P2X3 signaling in PNN and somatic tubulin accumulation are class-effects of PIs, we examined three additional compounds: (i) delanzomib, a peptide boronic acid like bortezomib that has been tested in clinical trials; (ii) epoxomicin, an epoxyketone like carfilzomib; and (iii) the peptide aldehyde MG-132. Delanzomib neither affected the viability nor the neurite growth in the PeriTox at test concentrations up to 10 μM (Fig. 7A). Pre-treatment of PNN (>DoD38) with concentrations as low as 5 nM lead to the attenuation of Ca^2+^-signaling upon P2X3 stimulation (Fig. 7B), while TRPV1 signaling was not impaired (Fig. S9B). As previously observed with bortezomib, a slight increase in resting cell [Ca^2+^]_i_ was detected at delanzomib concentrations associated with inhibition of P2X3 signaling (Fig. 7C). Thus, delanzomib, which inhibits the proteasome with a similar K_i_ as bortezomib [57], also showed here similar *in vitro* effects as the PIs studied earlier.

**Figure 7:**
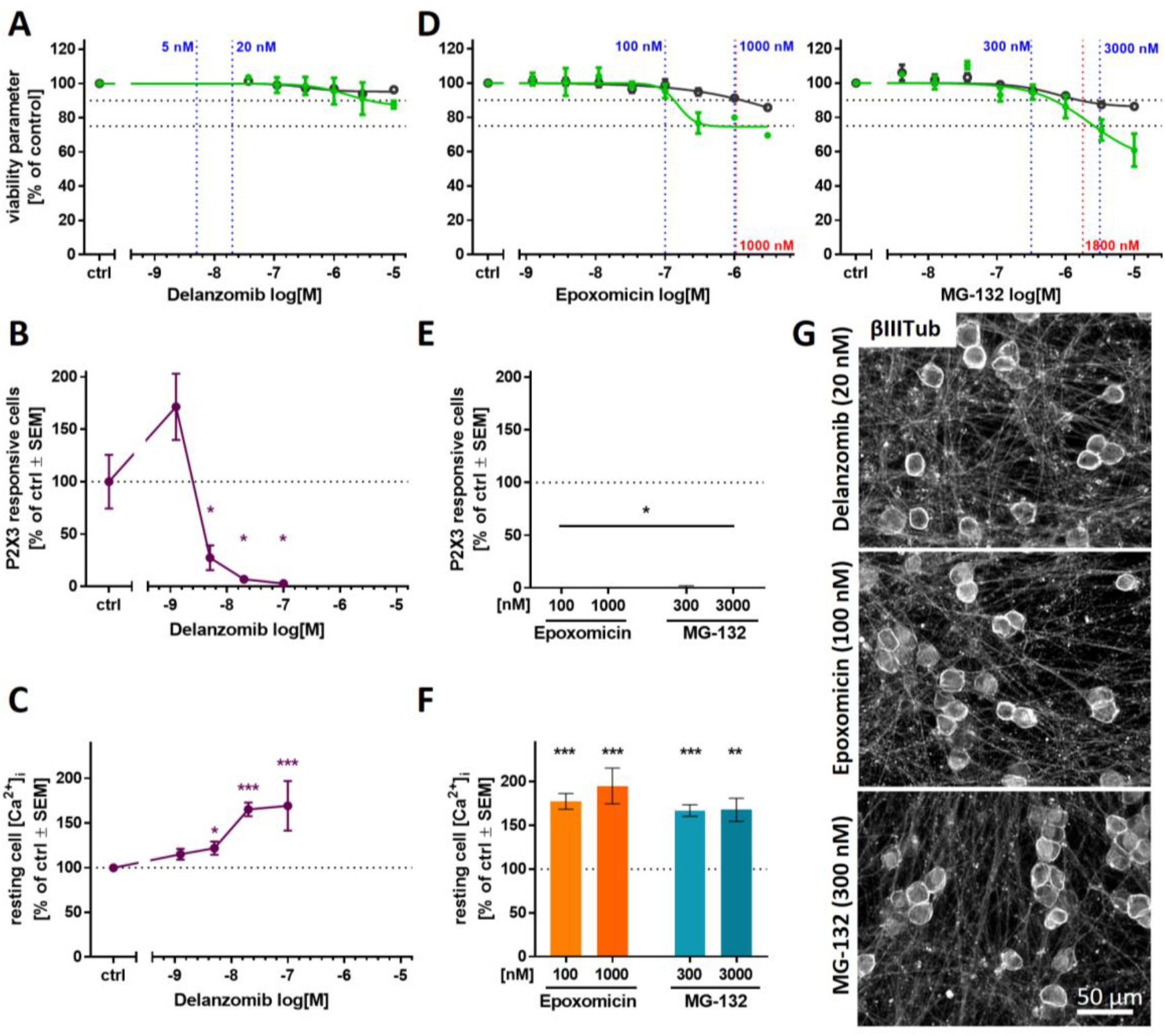
Attenuation of P2X3-signaling and microtubule reorganization as potential PI class-effects. The PIs delanzomib **(A-C,G)**, epoxomicin and MG-132 **(D-E,G)** representing different PI classes were investigated regarding their effects on various test endpoints. **(A,D)** The compounds’ effects on neurite area and viability were assessed in the standard PeriTox test. Horizontal dashed lines at 90% and 75% indicate the cytotoxicity threshold the neurite effect threshold, respectively. Vertical dashed lines indicate the lowest cytotoxicity-inducing concentration (red) and concentrations further used for Ca^2+^-imaging experiments (blue). **(B,C,E,F)** Sensory neurons (>DoD38) were pre-treated with the test compounds for 24 h before Ca^2+^-imaging experiments were performed. **(B,E)** The number of cells responding to stimulation with the P2X3-specific agonist α,β-methylene ATP (1 μM) was assessed. **(C,F)** Baseline fluorescence, indicating the resting cell intracellular [Ca^2+^]_i_ was quantified for whole sensory neuron cultures. **(A-F)** Data are given as % of untreated control cells and are means ± SEM of at least 3 biological replicates. * p < 0.05, ** p < 0.001, *** p < 0.0001. **(G)** After differentiation of >38 days, sensory neurons were exposed to the PIs for 24 h, fixed and stained for βIII-tubulin. Representative immunofluorescence images are shown. The scale bar is given in the images. Further details and quantification of cells with circular αIII-tubulin staining are given in figure S12.

Pre-screening of the experimental PIs epoxomicin and MG-132 in the PeriTox test revealed high cytotoxicity thresholds of ≥1,000 nM (Fig. 7D). For both compounds, test concentrations were chosen that did not alter any PeriTox test endpoint (100 nM epoxomicin and 300 nM MG-132). Pre-treatment of PNN to such conditions resulted in a complete blunting of P2X3 responses, accompanied by elevated [Ca^2+^]_i_ in resting cells (Fig. 7E,F). For all three PIs, we found that the neurite network remained intact upon exposure to P2X3-attenuating concentrations. As expected, we found that inhibition of the P2X3 responses again correlated with the emergence of sharp annular βIII-tubulin accumulations in the cell somata (Fig. 7G, S12A,B).

These results suggest that attenuation of P2X3 signaling and αIII-tubulin reorganization are indeed class-effects of PIs.

## Discussion & Conclusion

We have developed and documented here a robust differentiation protocol that yields human PNN useful to address various biomedical questions. The cultures generated in this way have many characteristics of nociceptors, and they can be used reproducibly after 6 weeks of differentiation (without cell detachment, with no signs of de-differentiation, and completely without any overgrowth by non-wanted cells) for single cell Ca^2+^-imaging of P2X3 receptors. The cells maintain their original network of individual somata, connected by long neurites. This is noteworthy, as many other culture protocols designed to yield peripheral neurons tend to generate cells that cluster together over time, and that make imaging of [Ca^2+^]_i_ in individual cells nearly impossible. These PNN allowed us to study very early adverse effects of PIs at clinically-relevant low nM concentrations [58, 59]. All five compounds investigated behaved similarly in that they induced a pronounced down-regulation of P2X3 responses and a clustering of tubulin to ring-like structures around the somata, at concentrations that were non-cytotoxic and that did not damage any of the neurite network features.

The PeriTox test is a well established *in vitro* screening assay using human iPSC-derived peripheral neurons to identify peripheral neurotoxicants [28]. It has been successfully used to identify environmental neurotoxicants by assessing their effects on the neurite structures [40–42]. However, for pre-clinical drug testing, an increased sensitivity in detecting the potential neurotoxicity of chemotherapeutics is desirable. To achieve this, we pursued different strategies:

First, we explored whether longer exposure times would decrease the toxicity threshold concentrations [60–62]. We found that prolonged exposure to toxicants increased the sensitivity for cytotoxicity, but the specificity for neurite effects was lost. This is in good agreement with the fact that in many neuronal cultures neurite damage is followed by general cell death or apoptosis, if given sufficient incubation time [63–65]. The sequence of neurite damage triggering cell death may be particularly pronounced in still differentiating iPSC-derived neurons, while *in vivo* matured neurons that are functionally integrated in regulatory circuits are known to separate the neurite pruning program from downstream death of the somata [66, 67].

As second approach, we explored whether functional changes in sensory receptors would allow for more sensitive readouts. Indeed, signaling through P2X3 receptors proved to be highly sensitive to proteasome inhibitors. The measurement of such responses required a new culture setup, using cells differentiated for ≥5 weeks. The detection of PI-induced effects by the new approach at ≥10-fold higher sensitivity than in the PeriTox test suggests that alterations at the functional level of signaling may often precede structural impairments.

Further examinations of timing aspects appear highly relevant. It would be interesting to learn whether P2X3 signaling remains a specifically altered endpoint upon prolonged exposure (48-72 h) to low nM concentrations, or whether specificity is lost, as already observed in the extended PeriTox test. Also repeated exposure scenarios are of interest, as they might model a possible accumulation of PIs in the DRG [58, 68].

Although we used the pronounced regulation of P2X3 here mainly as indicator of dysregulation, we wondered whether this may also play a pathophysiological role. Indeed, P2X3 is part of several complex pain regulation circuits. E.g., the acid sensing ion channel ASIC3, which is also involved in pain signaling, can lead to inhibition of P2X3 responses [69, 70]. Since bortezomib induces aerobic glycolysis and thus extracellular acidification, the above process may play a role in tissue [71]. Whether an interaction of P2X3 and ASIC3 is relevant in PNN needs to be clarified. Our results further show that P2X3 responses and intracellular baseline Ca^2+^ levels are de-regulated at identical toxicant concentrations. Thus, blunted P2X3 responses could be caused by or function as indicator of Ca^2+^ de-regulation [72, 73].

Coinciding with the functional effect of P2X3 attenuation, we detected a somatic accumulation of tubulin in PI-treated PNN. Our conclusion that tubulin accumulation is a PI class effect is further supported by a study on the PI lactacystin (not used here), which elicited the same pattern of tubulin re-organization into sharp rings [6]. Furthermore, somatic accumulation of cytoskeletal proteins upon PI treatment was reported also in mouse *in vivo* studies, suggesting that tubulin re-organization observed *in vitro* also occurs in animals [56]. It will be interesting to study a potential association of tubulin accumulation and changes in axonal transport. Since Ca^2+^ is also known to be a regulator of the cytoskeleton [74], de-regulation of [Ca^2+^]_i_ may be a common cause of P2X3 signaling impairments and morphological changes observed in PI-treated PNN.

When taxol was compared here to the class of PIs, we neither observed P2X3 inactivation, nor tubulin rings. Thus, different initial processes may be involved in the development of taxol peripheral neuropathies. Future experiments should test more classes of neuropathy-inducing cytostatics, such as platinum compounds or vinca alkaloids.

Overall, this study demonstrates the feasibility of developing target cell-specific test methods that are based on human cells. Using neuronal cultures other than peripheral neurons for research on chemotherapy-induced peripheral neuropathy can miss functional effects only detectable in the relevant target cells [28, 31]. Moreover, the use of high toxicant concentrations and of blunt endpoints (such as cell death) may make it very difficult to identify compounds that would attenuate the toxicity. We suggest that insights on specifically-impaired processes are important for the development of pharmacological countermeasures for peripheral neuropathies.

## Materials and Methods

### Differentiation of human iPSCs to peripheral neurons

We used the human iPSC lines mciPS (model no. SC301A-1; System Biosciences, Palo Alto, CA, USA), SBAD2 [75], Sigma iPSC0028 (Si28) (EPITHELIAL-1, #IPSC0028, Merck, Darmstadt, Germany) and the transgenic iPSC line Si28-NGN1. IPSC cultures were maintained under xeno-free conditions (see supplementary methods) [76].

The differentiation procedure for all iPSC lines is detailed in the supplementary methods (see also table S1). In brief, iPSCs were neuralized by dual SMAD inhibition followed by direction of the differentiation towards the sensory neuron fate using small molecule inhibitors [77]. After 9-12 days of differentiation, immature peripheral neurons were frozen in 90% fetal bovine serum (FBS) (Thermo Fisher Scientific, Waltham, MA, USA) and 10% dimethyl sulfoxide (DMSO; Merck, Darmstadt, Germany). After thawing, further maturation was driven by the growth factors glia-derived neurotrophic factor (GDNF, 25 ng/ml), brain-derived neurotrophic factor (BDNF, 12.5 ng/ml) and nerve growth factor (NGF, 25 ng/ml) (all from Bio-Techne, Minneapolis, MN, USA). For the differentiation of peripheral neurons with nociceptor features, doxycycline (2 μg/ml) exposure from DoD4’-9’ and DoD1-14 was integrated in the basis small molecule differentiation protocol, starting from Si28-NGN1 iPSC.

Peri.4U cells were provided by Axiogenesis (Cologne, Germany) and maintained according to the manufacturer’s protocol.

### Generation of the gene-edited iPSC line Si28-NGN1

Analogous to Boisvert et al. [43], an iPSC line with inducible NGN1-overexpression was created. A lentiviral sequence was designed to express the human NGN1 gene under control of a Tet-responsive element (TRE), which is dependant on the presence of Doxycycline (Dox). The expression of NGN1 was linked to the expression of turboRFP to monitor the induction. For selection, a hygromycin restistance gene was included. The vector and the principle of the fusion construct was published earlier [78]. The cell line was authenticated by short tandem repeat DNA typing and pluripotency was confirmed (data not shown) [79].

### Toxicity testing by assessment of neurite area and cell viability

Immature peripheral neurons were thawed and seeded at a density of 100,000 cells /cm^2^. For initial toxicity assessment, cells were left to attach for 1 h at 37°C, 5% CO_2_ followed by treatment with the test compounds. Cells were exposed to the compounds for 24 h, 48 h and 72 h and readout was performed on DoD1, 2 and 3, respectively.

For delayed toxicity assessment, cells were cultured until DoD4. On DoD4, test compounds were added to the cells together by performing a half medium exchange. Readout was performed after 72 h (on DoD7).

For the readout, neurons were stained with 1 μg/ml HOECHST-33342 (H-33342) and 1 μM calcein-AM (both from Merck, Darmstadt, Germany) one hour prior to the imaging. After incubation for 1 h at 37°C, 5% CO_2_, images were acquired automatically using an ArrayScan VTI HCS microscope (Thermo Fisher Scientific, Waltham, MA, USA). Images were analysed for neurite area and cell viability as previsouly described [80].

### Immunofluorescence staining

Protein expression was assessed qualitatively via immunofluorescence staining and microscopy. All samples were prepared, and analyzed exactly as described before [39, 81], using antibodies as detailed in supplementary methods.

The quantification of somata with βIII-tubulin circles was performed manually by independent observers. Criteria for the quantification were sharply defined, intense, and circular βIII-tubulin staining with presumably sub-membraneous location covering at least 50% of a full circle.

To quantify the nuclear area, images of H-33342 stainings were converted into binary images and the area of randomly chosen H-33342-objects was measured using the Fiji software.

### Transcriptome data generation and analysis

Sample lysates were prepared by medium removal, followed by a wash with 50 μl of phosphate buffered saline (PBS) (Thermo Fisher Scientific, Waltham, MA, USA) and instant addition of 33 μl 1x Biospyder lysis buffer (BioSpyder Tech., Glasgow, UK) [39, 82]. After incubation at RT for 10 minutes, the sample plates were stored at −80°C up to the time of dry ice shipping to Bioclavis (BioSpyder Tech., Glasgow, UK). Measurement of the whole transcriptome set was performed via the TempO-Seq targeted sequencing technology [83]. The gene set analyzed, and the read data are detailed in supplementary file 2 (organized as Excel workbook). For data processing, the R package DESeq2 (v1.32.0) was used for quality control and normalization [84]. Further details on data analysis are given in the supplementary methods.

### Measurement of changes in intracellular Ca^2+^ concentration [Ca^2+^]_i_

Sensory neurons were cultured in 96-well plates for at least 35 days after thawing. One day before Ca^2+^-imaging experiments were performed, cells were optionally pre-treated with the test compounds. One hour before the experiment, cells were loaded with the Ca^2+^-indicator Fluo-4 (Thermo Fisher Scientific, Waltham, MA, USA). Pre-treatment with antagonists was performed together with Fluo-4 loading. Monitoring of [Ca^2+^]_i_ was performed using an ArrayScan VTI HCS microscope equipped with an automated pipettor and an incubation chamber providing an atmosphere with 5% CO_2_ and 37°C. Cells were imaged as fast as possible for 45 s. Test compounds were automatically applied after baseline recording (10 s). In a standard experiment with 4 stimuli applied to one well (e.g., negative control, P2X3 agonist, TRPV1 agonist, KCl), the cells were imaged 4 times for 45 s with one stimulus applied at a time.

The images were exported as *.avi video files and analysed with the CaFFEE software [85]. In brief, the time point of peak fluorescence was identified. Fluorescence data for the ground state (F_0_) and for the peak time point (F_1_) were assessed automatically for all cells. The difference between the two fluorescence levels, ΔF=F_1_-F_0_, was used for further data processing [39]. The noise level-based threshold (mean(ΔF) + 3x SD(ΔF)) of each well was determined by the application of a negative control stimulus (Hanks’ Balanced Salt Solution (HBSS)), with an upper threshold-limit set to ΔF=18. According to this threshold, cells were defined as reactive (ΔF_stimulus_>threshold) or non-reactive (ΔF_stimulus_ ≤threshold).

### Statistics

If not stated otherwise, experiments were performed on 3 or more independent cell preparations (here called biological replicates). In each cell preparation at least three different wells (here called technical replicates) were measured.

Information concerning descriptive statistics and experimental variability is included in the figure legends or the figures themselves. GraphPad Prism 7 software (Version 7.04, Graphpad Software, Inc, San Diego, USA) was used for significance testing and data display. Data were evaluated by ANOVA plus appropriate post-hoc testing method or by t-test for binary comparisons. *p*-values < 0.05 were regarded as statistically significant.

## Acknowledgements

This work was supported by CEFIC, the BMBF, EFSA, and the DK-EPA (MST-667-00205). It has received funding from the European Union’s Horizon 2020 research and innovation program under grant agreements No. 681002 (EU-ToxRisk), No. 964537 (RISK-HUNT3R), No. 964518 (ToxFree) and No. 825759 (ENDpoiNTs). We are grateful to M. Kapitza for invaluable experimental support.

## Disclosure of Potential Conflicts of Interest

The authors declare no conflict of interest.

## Abbreviations

[Ca^2+^]_i_: intracellular Ca^2+^-concentration
α,β-meATP: α,β-methylene ATP
βIIITub: βIII-tubulin
AraC: cytarabine
ATP: adenosine triphosphate
BIPN: bortezomib-induced peripheral neuropathy
BDNF: brain-derived neurotrophic factor
BTZ: bortezomib
CFZ: carfilzomib
CNS: central nervous system
CPM: counts per million
DMSO: dimethyl sulfoxide
DRG: dorsal root ganglion
FBS: fetal bovine serum
FGF: fibroblast growth factor
GDNF: glia-derived neurotrophic factor
HBSS: Hanks’ Balanced Salt Solution
iPSC: induced pluripotent stem cell
NGF: nerve growth factor
PI: proteasome inhibitor
PNN: peripheral neurons with nociceptor features
PNS: peripheral nervous system
PRPH: peripherin
RT-qPCR: quantitative reverse transcriptase polymerase chain reaction
ROCKi: ROCK inhibitor
TRP: transient receptor potential

## Notes

### Competing Interest Statement

The authors have declared no competing interest.

