## Supplementary File 1 for "Specific attenuation of purinergic signaling during bortezomib-induced peripheral neuropathy"

**Table of Contents**


| | | |
|---|---|---|
| | Page2 (P2) | Supplementary Methods |
| Tab. S1 | P4 | Schedule for differentiation media and supplementing factors |
| Tab. S2 | P5 | Primary antibodies used in this study |
| Tab. S3 | P5 | Agonists and antagonists used in $Ca^{2+}$-imaging experiment |
| Fig. S1 | P6 | Characterization of iPSC-derived early-stage sensory neurons |
| Fig. S2 | P7 | Exploration of toxicant responses (in the PeriTox test) of neurons generated and offered by a commercial supplier |
| Fig. S3 | P8 | Time-dependency of compound effects assessed in the extended PeriTox test |
| Fig. S4 | P9 | Variation of exposure time point and duration to assess compound toxicity |
| Fig. S5 | P10 | Characterization of peripheral neurons derived by NGN1-overexpression |
| Fig. S6 | P11 | Expression of functional TRPV1 channels |
| Fig. S7 | P12 | Time dependent transcriptome profiling of PNN |
| Fig. S8 | P13 | $[Ca^{2+}]_i$-dependent fluorescence signals of single cells |
| Fig. S9 | P14 | Membrane depolarization and TRPV1 responses are unaltered upon toxicant exposure |
| Fig. S10 | P15 | Reorganization of the cytoskeletal structure upon PI exposure |
| Fig. S11 | P17 | Quantification of the nuclear area as cell viability indicator |
| Fig. S12 | P18 | Reorganization of the microtubule structure as a general PI effect |
| | P20 | Supplementary references |

**Supplementary Methods**

***Maintenance of induced pluripotent stem cells (iPSCs)***

Maintenance of all iPSC lines used in this study was performed on human Laminin-521 (BioLamina, Sundbyger, Sweden) coating in essential 8 (E8) medium (Dulbecco's modified Eagle's medium/F12 [DMEM/F12] supplemented with 15 mM Hepes [Thermo Fisher Scientific, Waltham, MA, USA], 10 µg/ml holo-transferrin, 20 µg/ml insulin, 16 mg/ml L-ascorbic-acid, 0.7 mg/ml sodium selenite [all from Merck, Darmstadt, Germany], 100 ng/ml bFGF [Thermo Fisher Scientific, Waltham, MA, USA], 1.74 ng/ml TGFb [Bio-Techne, Minneapolis, MN, USA]) essentially as described [1]. Passaging of the iPSCs was performed every 7 days. Cells were incubated with EDTA (Thermo Fisher Scientific, Waltham, MA, USA) for 2 min (37°C, 5% $CO_2$) to detach the cells, so that clumps remain (no single cell suspension). IPSCs were washed off the plate with DMEM/F12. Cells were re-seeded in E8 medium on freshly coated plates in a final dilution of 1:40-60.

***Differentiation of sensory neurons from iPSC***

The iPSCs were prepared for neural differentiation on day of differentiation minus 2 (DoD-2) by replating in a single cell suspension (90,000 cells/cm$^2$) onto Matrigel$^{TM}$ (Corning, Glendale, AZ, USA) coated 6-well plates in E8 medium supplemented with 10 µM Rock inhibitor (Y-27632 [Bio-Techne, Minneapolis, MN, USA]).

On DoD0', E8 was replaced by neural differentiation medium KSR (knock out DMEM with 15% serum replacement, 1 x Glutamax, 1 x nonessential amino acids, and 50 µM β-mercaptoethanol [all from Thermo Fisher Scientific, Waltham, MA, USA]) and the combination of five small molecule pathway inhibitors as described in detail in table S1. From DoD0'-5', Noggin and SB-431642 (both from Bio-Techne, Minneapolis, MN, USA) (± dorsomorphine hydrochloride [Dorso; Merck, Darmstadt, Germany]) were added, and CHIR99021 (Axon Medchem, Groningen, Netherlands), SU5402 (Bio-Techne, Minneapolis, MN, USA) and DAPT (γ-Secretase inhibitor IX; Merck, Darmstadt, Germany) were added on DoD2'-9'. From DoD4' onwards, KSR medium was gradually replaced by N2-S medium (DMEM/F12, 1 x GlutaMax [both from Thermo Fisher Scientific, Waltham, MA, USA], 0.1 mg/ml apotransferrin, 1.55 mg/ml glucose, 25 µg/ml insulin, 20 nM progesterone, 100 µM putrescine and 30 nM selenium [all from Merck, Darmstadt, Germany]) in 25% increments. On DoD9' or DoD12' (depending on the iPSC line used, see table S1), the cells were cryopreserved in 90% fetal bovine serum (FBS) (Thermo Fisher Scientific, Waltham, MA, USA) and 10% dimethyl sulfoxide (DMSO; Merck, Darmstadt, Germany).

After thawing of the pre-differentiated cells, sensory neuron precursors were cultured in 25% KSR and 75% N2-S supplemented with CHIR99021 (1.5 µM), SU5402 (5 µM) and DAPT (5 µM). Cells were seeded at a density of 100,000 cells/cm$^2$ on Matrigel$^{TM}$ coated plates. For further differentiation and maturation, half of the medium was changed on DoD1 and DoD2. With the fresh culture medium on DoD2, Matrigel was added to the cells at a final dilution of 1:80. On DoD3, medium was changed to N2-S medium supplemented with 12.5 ng/ml brain-derived neurotrophic factor (BDNF), 25 ng/ml glia-derived neurotrophic factor (GDNF) and 25 ng/ml nerve growth factor (NGF) (all from Bio-Techne, Minneapolis, MN, USA) and 2 µM cytarabin (AraC; Merck, Darmstadt, Germany). Half of the medium was changed on DoD4, 7 and 10 with further Matrigel addition on DoD10. On DoD14, medium was changed to maturation medium (N2-S supplemented with BDNF [12.5 ng/ml], GDNF and NGF [both 25 ng/ml]). Half medium exchanges were performed every three to four days. Matrigel was diluted in the culture medium at a final dilution of 1:80 every 10 days.

### *Assessment of neurite area and cell viability*

Cells stained with calcein-AM and HOECHST-33342 (H-33342) (both from Merck, Darmstadt, Germany) were imaged automatically using an ArrayScan VTI HCS microscope (Thermo Fisher Scientific, Waltham, MA, USA). Images were analysed for neurite area and cell viability by an automated algorithm. H-33342 staining was used to identify the cell nuclei. The nuclear area was enlarged to define the somatic area, which was then substracted from the calcein stain. The remaining calcein-positive pixels were quantified to derive the neurite area. Data on cell viability was derived from the same images by checking each H-33342 stained object (cell) for a double stain with calcein. Double-positive cells were classified as viable, calcein-negative cells as dead [2].

### *Immunofluorescence staining and microscopy*

Neurons, grown on glass coverslips coated with Matrigel™, were fixed with 4% paraformaldehyde at 4°C over night. All further steps were performed at room temperature. Paraformaldehyde was taken off and cells were washed (~1 min) with phosphate buffered saline (PBS) followed by permeabilization with 0.6% Triton X-100 in PBS for 7 min. Coverslips were washed (~1 min) with PBS and blocked for 1 h in PBS containing 5% FCS and 0.1% Triton X-100. Primary antibodies (see Tab. S2) were diluted in fresh blocking solution and applied for 1 h. Residual free primary antibodies are then washed off with PBS. Secondary antibodies and H-33342 are diluted in blocking solution and applied on the coverslips for 30 min. After washing with PBS, coverslips were placed upside-down on mounting medium on microscope slides.

### *Transcriptome data analysis*

For data processing, the R package DESeq2 (v1.32.0) was used for quality control, normalization and differential gene expression (DGE) analysis [3]. The raw probe counts were normalized to total sample counts per million (CPM). No library size threshold was used; samples with replicate correlation (Pearson R) to group average below 0.8 were removed from the analysis. Prior to the DGE, the low-count genes (less than 3 samples above 5 CPM) were discarded. The Wald statistics test was used for significance evaluation of each differentiation stage against DoD1 or DoD7. Selection of the most significant differentially expressed genes (DEG) was done using a (Benjamini-Hochberg) p-adjusted maximum threshold of 0.05 and a $log_2$(fold change) minimum threshold of 1.

***Supplementary table 1: Schedule for differentiation media and supplementing factors***

**X**: applies for all iPSC lines
**X**: applies for mc-iPSC
**X**: applies for SBAD2
**X**: applies for Si28-NGN1

| **DoD** | **Medium** | | **Supplements** | | | | | | |
|---|---|---|---|---|---|---|---|---|---|
| | KSR | N2S | Noggin (17.5 ng/ml) | SB431542 (10 µM) | Dorso (600 nM) | CHIR99021 (1.5 µM) | SU5402 (5 µM) | DAPT (5 µM) | Dox (2 ng/ml) |
| 0′ | **X** | | **X** | **X** | **X** | | | | |
| 1′ | **X** | | **X** | **X** | **X** | | | | |
| 2′ | **X** | | **X** | **X** | **X** | **X** | **X** | **X** | |
| 3′ | **X** | | **X** | **X** | **X** | **X** | **X** | **X** | |
| 4′ | **X** (75%) | **X** (25%) | **X** | **X** | **X** | **X** | **X** | **X** | **X** |
| 5′ | **X** (50%) | **X** (50%) | | | | **X** | **X** | **X** | **X** |
| 6′ | **X** (50%) | **X** (50%) | | | | **X** | **X** | **X** | **X** |
| 7′ | **X** (25%) | **X** (75%) | | | | **X** | **X** | **X** | **X** |
| 8′ | **X** (25%) | **X** (75%) | | | | **X** | **X** | **X** | **X** |
| 9′ | **X** (25%) | **X** (75%) | | | | **X** | **X** | **X** | |
| 10′ | **X** (25%) | **X** (75%) | | | | **X** | **X** | **X** | |
| 11′ | **X** (25%) | **X** (75%) | | | | **X** | **X** | **X** | |

***Supplementary table 2: Primary antibodies used in this study***

| **Target** | **Isotype** | **Dilution** | **Supplier** | **Catalogue number** |
|---|---|---|---|---|
| βIII-tubulin (pol.) | mouse IgG1 | 1:1000 | BioLegend | 921001 |
| Peripherin | mouse IgG2a | 1:200 | Santa Cruz | sc-377093 |
| Ki67 (PE) | mouse IgG1 | 1:500 | BD Pharmingen | 556027 |
| ISL1 | rabbit | 1:200 | Abcam | ab109517 |
| BRN3A | rabbit | 1:200 | Merck Millipore | 5945 |
| P2X3 | rabbit | 1:200 | Novus | NB100-1654 |
| F-actin | Phalloidin-555 | 1:500 | Molecular Probes | 8953S |

***Supplementary table 3: Agonists and antagonists used in $Ca^{2+}$-imaging experiments***

| **Compound** | **Solvent** | **Concentration [μM]** | **Supplier** | **Catalogue number** |
|---|---|---|---|---|
| α,β-methylene ATP | water | 1 | Cayman Chemical | 10008956 |
| AF-353 | DMSO | 0.1 | Cayman Chemical | 23034 |
| capsaicin | DMSO | 1 | Cayman Chemical | 92350 |
| capsazepine | DMSO | 10 | Cayman Chemical | 10007518 |
| KCl | water | 40,000 | Merck | P9541 |

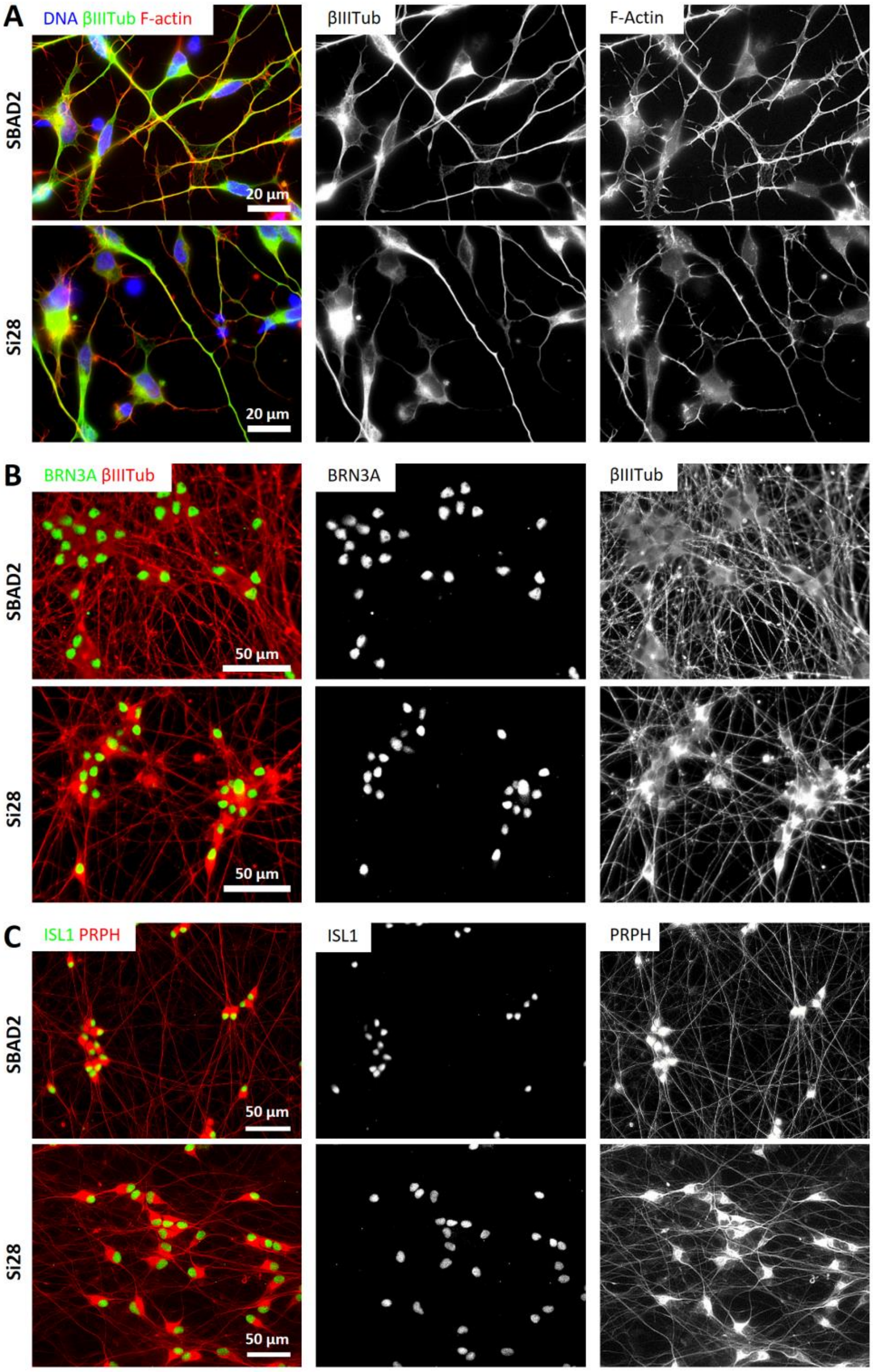


**Fig. S1**
**Holzer et al. 2022**

**Supplementary figure 1: Characterization of iPSC-derived early-stage sensory neurons**

Neurons were differentiated from the iPSC-lines SBAD2 and Si28 according to the standard protocol . **(A)** Cells were fixed on DoD1 after thawing and stained for the pan-neuronal marker βIII-tubulin (βIIITub), and for F-actin. Growth cones with filopodia are evident on the growing neurites. **(B,C)** Cells were fixed on DoD7 and stained for βIIITub and the peripheral neuron transcription factors BRN3A **(B)** or Islet-1 and the intermediate filament peripherin (PRPH) **(C)**. Images of different fluorescent channels are shown to visualize the staining by individual antibodies. The composite images of the stainings are color coded according to the detail images (also see figure 1A). Scale bars are given in the images. All images are representative of typical cultures, but structures have not been quantified.

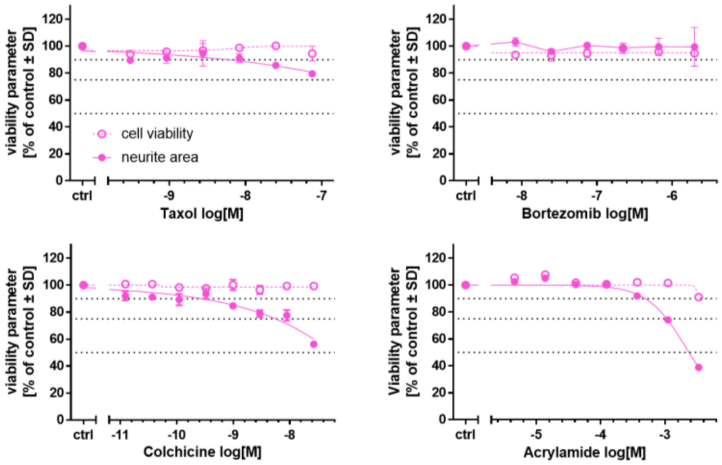


**Fig. S2**
**Holzer et al. 2022**

**Supplementary figure 2: Exploration of toxicant responses (in the PeriTox test) of neurons generated and offered by a commercial source.**

The Peri.4U cell preparation, which had been generated to provide a commercial source of peripheral neurons, was used in the PeriTox test. Cells were exposed to the positive control compounds taxol, bortezomib, colchicine and acrylamide, which are known for their (peripheral) neurotoxic potential. Effects on neurite area (solid symbols and lines) and viability (open symbols, dashed lines) are shown. Data are means ± SD of 2 biological replicates.

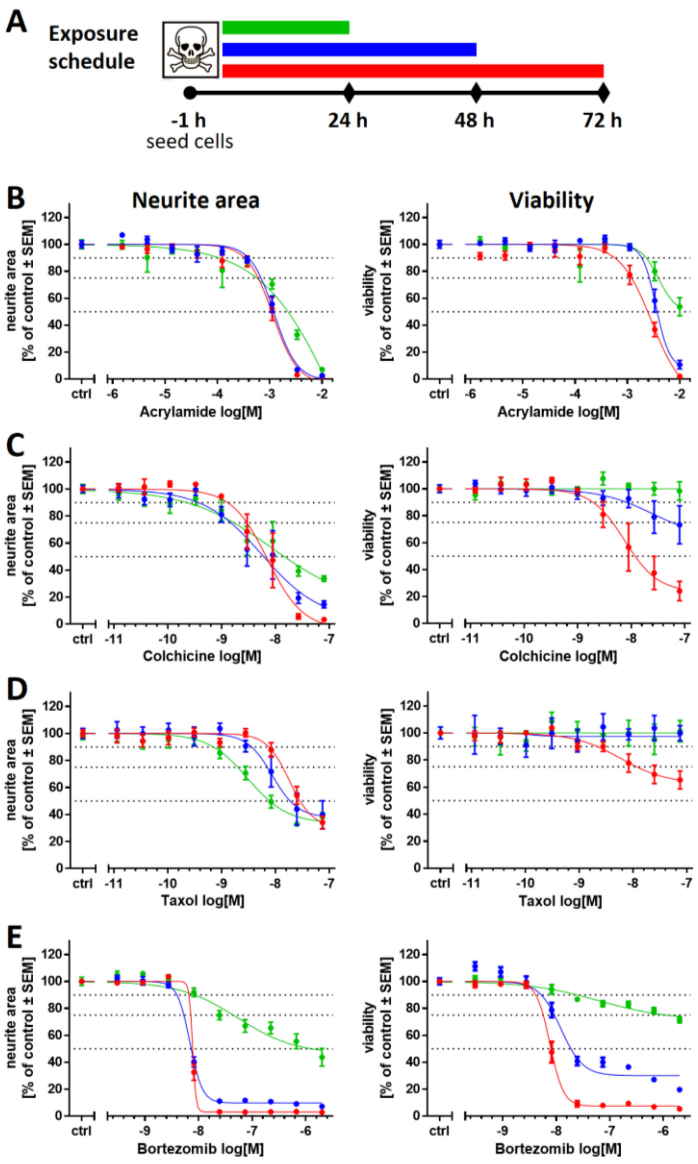


**Fig. S3**
**Holzer et al. 2022**

**Supplementary figure 3: Time-dependency of compound effects assessed in the extended PeriTox test**

Compound-induced effects on the neurite area and cell viability were assessed in the extended PeriTox test. (**A**) Exposure schedules for the extended PeriTox test are depicted. One hour after plating of the peripheral neuron precursors, cells were exposed to the test compounds for 24 h (green), 48 h (blue) and 72 h (red). **(B-E)** Effects on neurite area (left) and viability (right) are shown for all three exposure times for the positive controls acrylamide **(B)**, colchicine **(C)**, taxol **(D)**, and bortezomib **(E)**. Data sets are colour coded to the respective exposure schedule in **(A)**. Data are means ± SEM of 2-3 biological replicates.

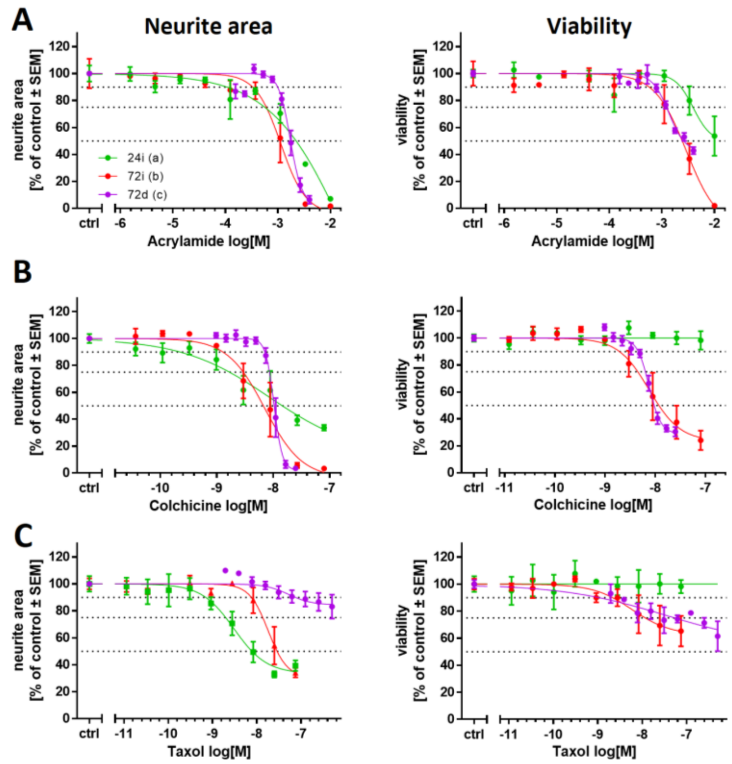


**Fig. S4**
**Holzer et al. 2022**

**Supplementary figure 4: Variation of exposure time point and duration to assess compound toxicity**

The positive control compounds acrylamide **(A)**, colchicine **(B)** and taxol **(C)** were tested regarding their effects on immature peripheral neurons. Cells were exposed to the test compounds according to the exposure scenarios depicted in figure 2A: (a) immediately after thawing (1 h) for 24 h (green, 24i) and (b) 72 h (red, 72i), or (c) at a delayed time point, starting on DoD4, for 72h (purple, 72d). Compound effects on neurite area (left) and viability (right) are shown. Data are means ± SEM of 3 independent experiments.

.

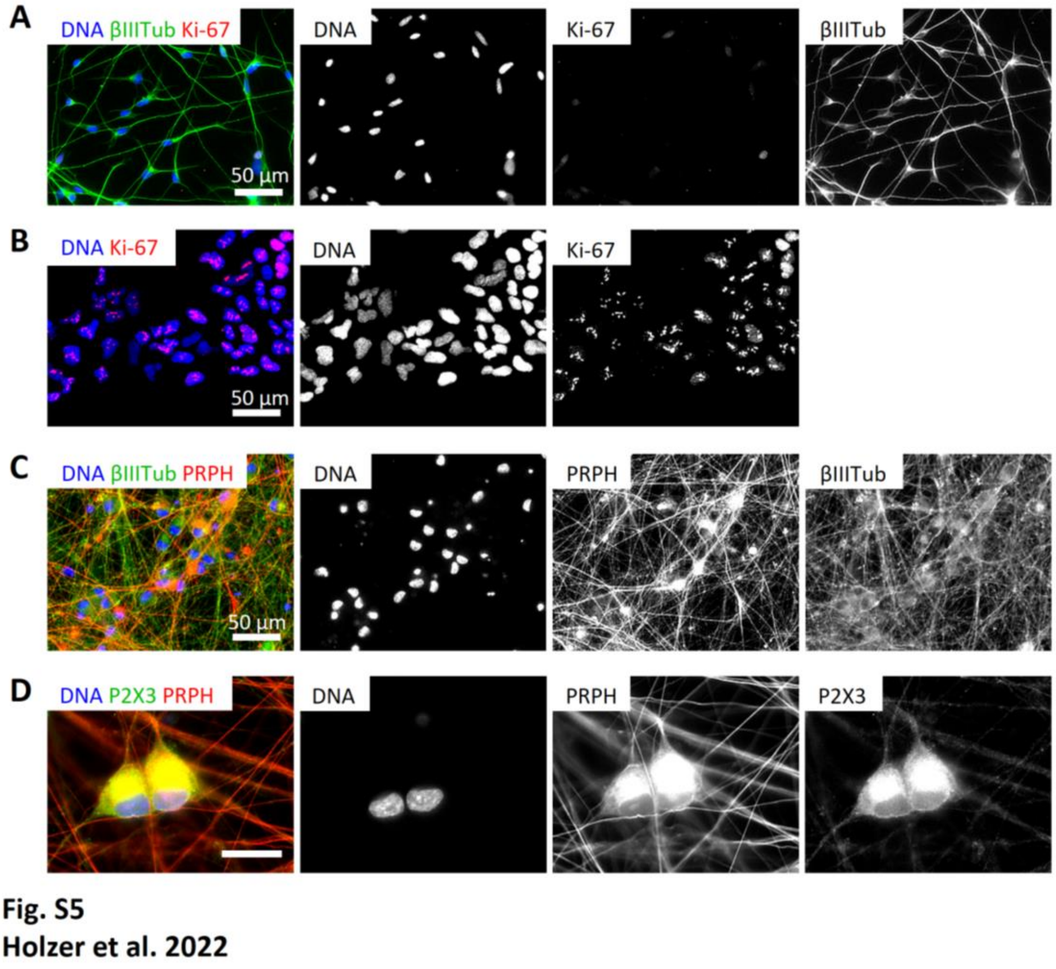


**Supplementary figure 5: Characterization of peripheral neurons derived by NGN1-overexpression**

**(A)** Peripheral neurons differentiated from Si28-NGN1 iPSCs were fixed on DoD1 and stained for βIII-tubulin (βIIITub) and the proliferation marker Ki-67. **(B)** As a positive control for proliferating cells, neuroepithelial precursor cells [4] were fixed and stained for Ki-67, exhibiting the characteristic pattern of Ki-67-staining in the nucleus, thus verifying the functioning of the Ki-67-antibody. **(C,D)** Neurons were differentiated for 42 days after thawing and fixed. Immunofluorescence images show an extensive neurite network, stained positive for βIIITub and peripherin (PRPH) **(C)**, and the expression of P2X3 receptors in PRPH-positive cells **(D)**. Images of the single stainings are shown. The composite images of the stainings are color coded according to the detail images (**(A,C,D)** see also figure 3B). Scale bars are given in the images.

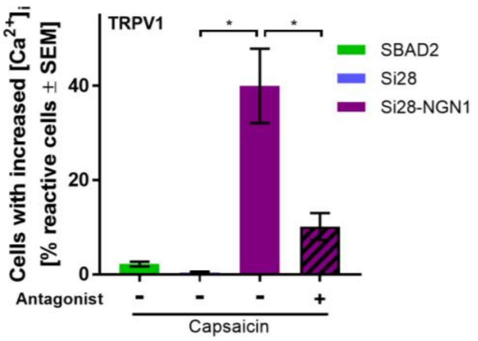


**Fig. S6**
**Holzer et al. 2022**

**Supplementary figure 6: Expression of functional TRPV1 channels**

Peripheral neurons derived from the iPSC lines SBAD2 (green), Si28 (blue) and Si28-NGN1 (purple) were differentiated for >38 days and used for $Ca^{2+}$-imaging experiments. The TRPV1-specific agonist capsaicin [1 µM] was used to determine the expression of functional TRPV1 receptors. To confirm signaling through TRPV1, cells were pre-treated (1 h) with capsazepine, an antagonist of TRPV1, before stimulation with capsaicin. Data are means ± SEM of at least 3 biological replicates.

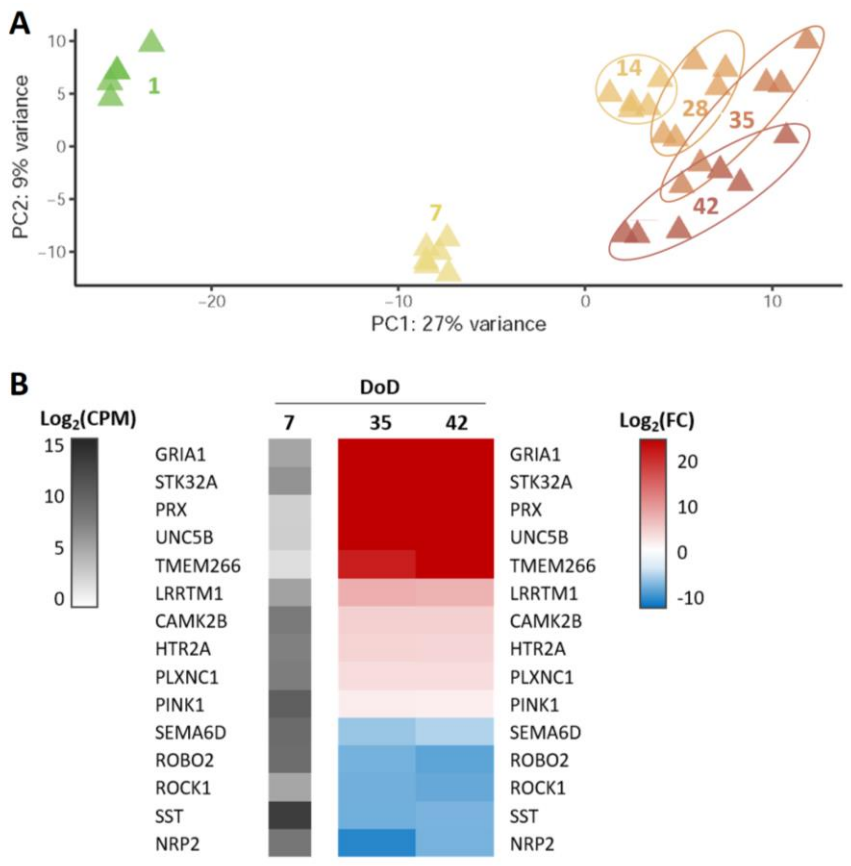


**Fig. S7**
**Holzer et al. 2022**

**Supplementary figure 7: Time-dependent transcriptome profiling of PNN**

**(A)** PNN were grown from DoD1 - DoD42 and mRNA of different maturation stages was used for whole transcriptome analysis by TempO-Seq (full data in Supplementary file1, complementary data in figure 4). A PCA was performed for all of the >19,000 measured genes. In the two-dimensional PCA display, six maturation stages (day of differentiation (DoD) 1, 7, 14, 28, 35 and 42) of PNN are color-coded according to their DoD. Data points are derived from three independent differentiations, and all individual samples analyzed are depicted. **(B)** To gain insight into gene expression changes that occur during late differentiation, DoD35 and DoD42 samples were analyzed for differentially expressed genes (DEGs) *versus* DoD7. Examples of significant DEGs (adjusted *p*-value ≤ 0.05) with a fold change (FC) >4 were assembled. The left column shows the absolute expression levels of selected significant DEGs on DoD7 in counts of the corresponding gene per 1 million reads (CPM). The data for DoD35/DoD42 show the fold change (FC) of the expression levels *versus* DoD7. The color scales use $\log_2$ units (see supplementary files for complete data sets).

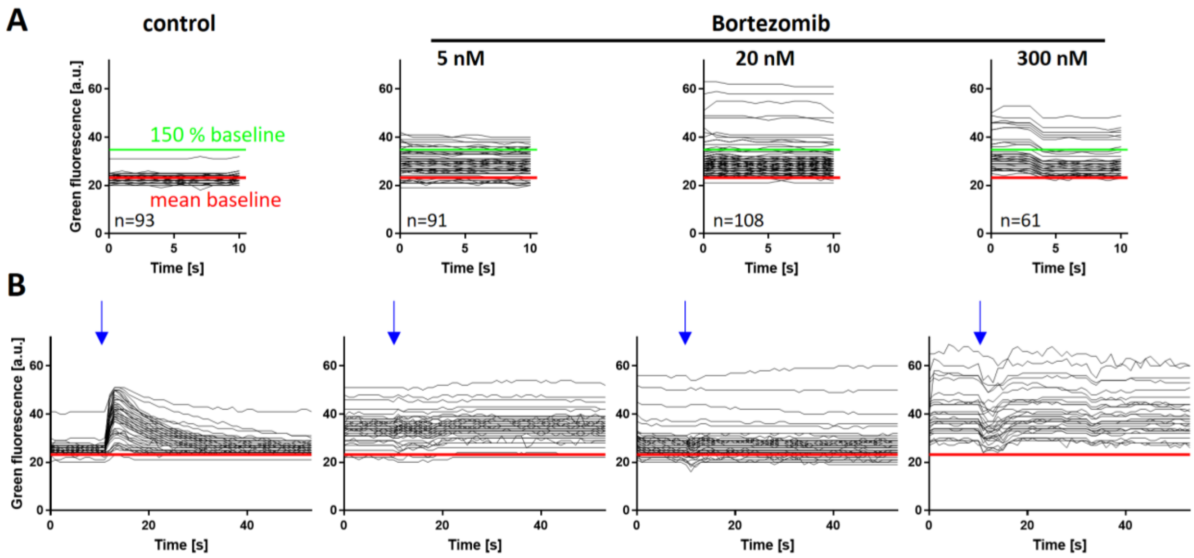


**Fig. S8**
**Holzer et al. 2022**

**Supplementary figure 8: [$Ca^{2+}$]$_i$-dependent fluorescence signals of single cells**

Sensory neurons (>DoD38) were cultured under control conditions or exposed to bortezomib for 24 h, before $Ca^{2+}$-imaging was performed. $Ca^{2+}$-measurements depicted in **(A)** and **(B)** were executed with the same cells. **(A)** Baseline fluorescence of single sensory neurons recorded during the first 10 seconds of a $Ca^{2+}$-measurement, immediately before the first stimulus application. Fluorescence traces of one whole neuronal culture per pre-treatment condition with 61-108 single cells each are shown. Red horizontal lines indicate the mean baseline fluorescence of control cells. Green lines indicate the threshold of 150% control baseline fluorescence used for quantification of cells with increased resting cell [$Ca^{2+}$]$_i$ (also see figure S13). **(B)** Time-dependent fluorescence traces of single cells during experiments using the P2X3-specific agonist α,β-methylene ATP (addition to the cells is indicated by blue arrows). For clarity, only 50% of the cells depicted in **(A)** are shown (randomly chosen by software). Red horizontal lines correspond to the mean baseline fluorescence of control cells determined in **(A)**.

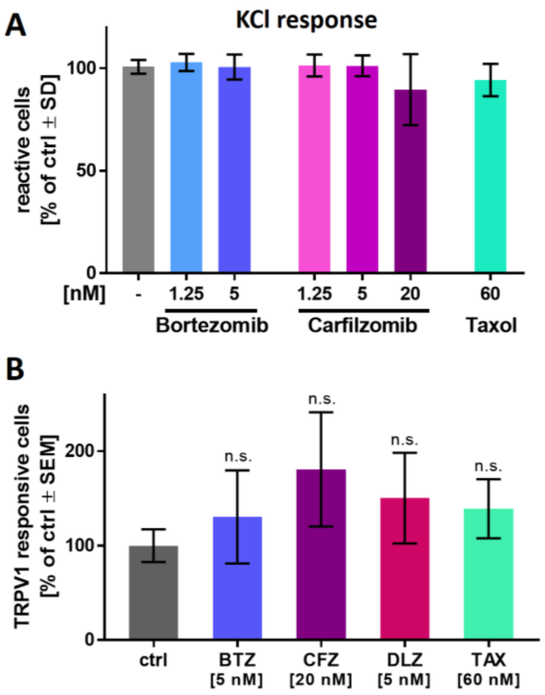


**Fig. S9**
**Holzer et al. 2022**

**Supplementary figure 9: Membrane depolarization and TRPV1 responses are unaltered upon toxicant exposure**

Sensory neurons were differentiated for >38 days and used for $Ca^{2+}$-imaging experiments. Cells were pre-treated with the proteasome inhibitors bortezomib (BTZ), carfilzomib (CZF) and delanzomib (DLZ), or taxol for 24 h, before $Ca^{2+}$-measurements were performed. **(A)** Sensory neurons, which were first exposed to a P2X3-specific stimulus (see figure 5B,C), were subsequently subjected to an increased concentration of KCl [50 mM], which induces membrane depolarization in functional neuronal cells. Responsive cells were quantified. **(B)** Cells were stimulated with capsaicin [1 µM] and the number of responsive cells was assessed. Data are given as % of untreated control cells and are means ± SEM of at least 3 independent experiments. n.s., not significant

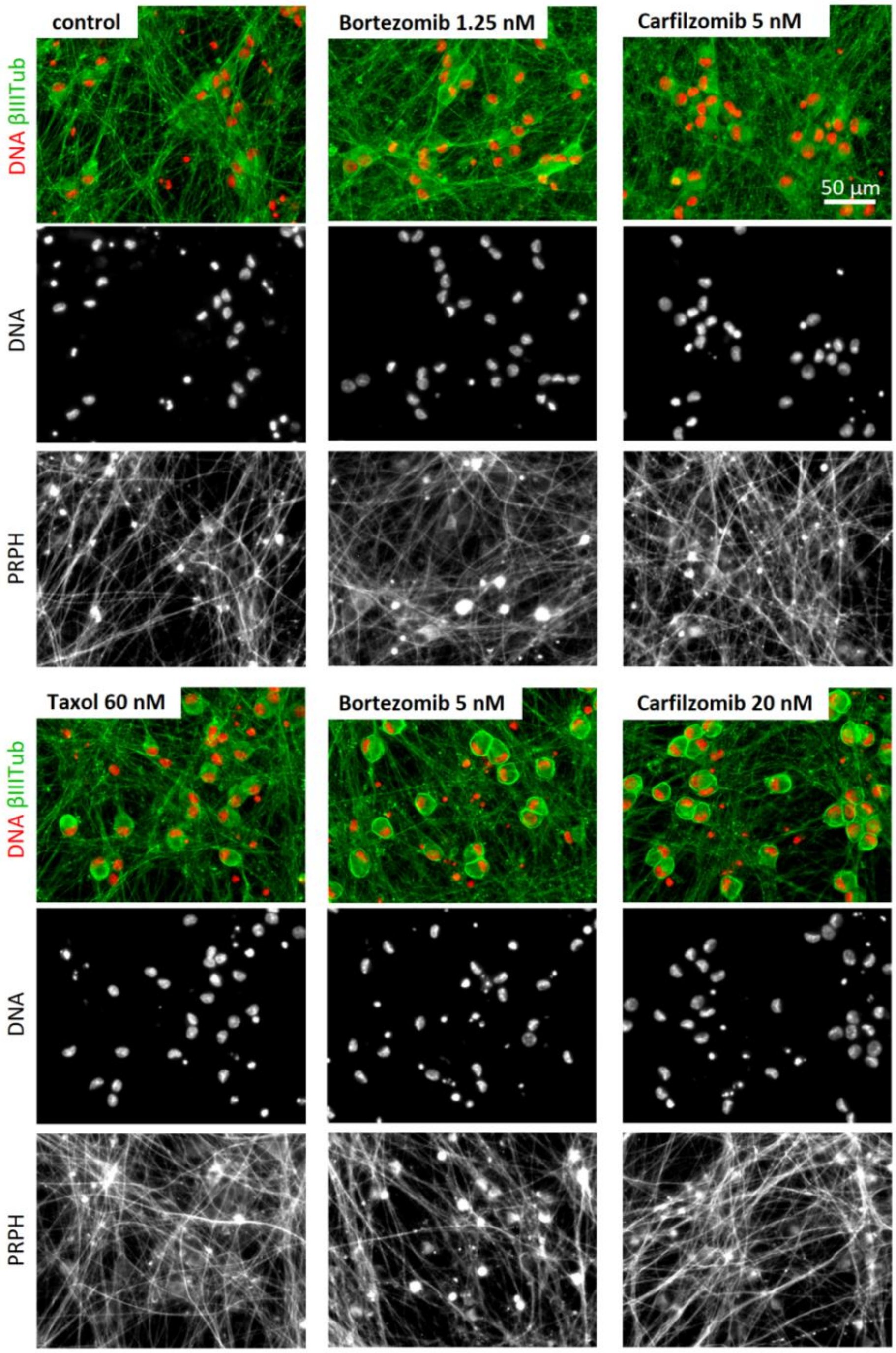


**Fig. S10**
**Holzer et al. 2022**

**Supplementary figure 10: Reorganization of the cytoskeletal structure upon PI exposure**

Sensory neurons were differentiated for at least 38 days after thawing and exposed to bortezomib, carfilzomib or taxol for 24 h before fixation. Representative immunofluorescence images are shown (see also figure 6A). Composite images show stainings of nuclei (DNA, red) and βIII-tubulin (βIIITub, green), revealing aberrant staining for βIIITub at high concentrations of bortezomib or carfilzomib. Single staining images shown for (i) the nuclei (DNA) and (ii) the corresponding peripherin (PRPH) staining (for clarity not included in the composite images). Peripherin staining of PI-treated cells did not exhibit βIIITub-like reorganization to rings, but rather a clustering in the soma (outside the nucleus). The scale bar is given in the images.

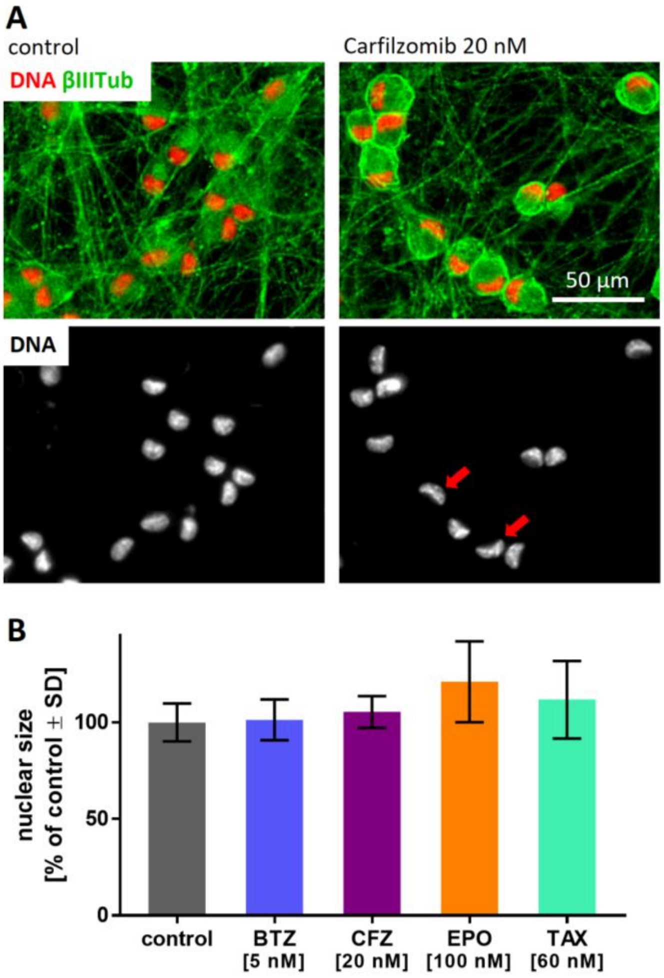


Fig. S11
Holzer et al. 2022

**Supplementary figure 11: Quantifiaction of the nuclear area as cell viability indicator**

**(A)** Sensory neurons were fixed on DoD42 upon exposure to control condition (left) or carfilzomib (20 nM, 24 h) (right). Immunofluorescence images of cells stained for βIII-tubulin (βIIITub) and DNA (using H33342) are shown. DNA staining revealed a deformation of the nuclei in carfilzomib-treated cells, red arrows indicate example nuclei in the single staining image. Images are color coded and the scale bar is given in the images. **(B)** Quantification of the nuclear area of cells cultured under control condition or exposed to bortezomib (BTZ), carfilzomib (CFZ), epoxomicin (EPO) or taxol (TAX) for 24 h at the indicated concentrations. Data are means ± SD of 25 randomly chosen nuclei per condition.

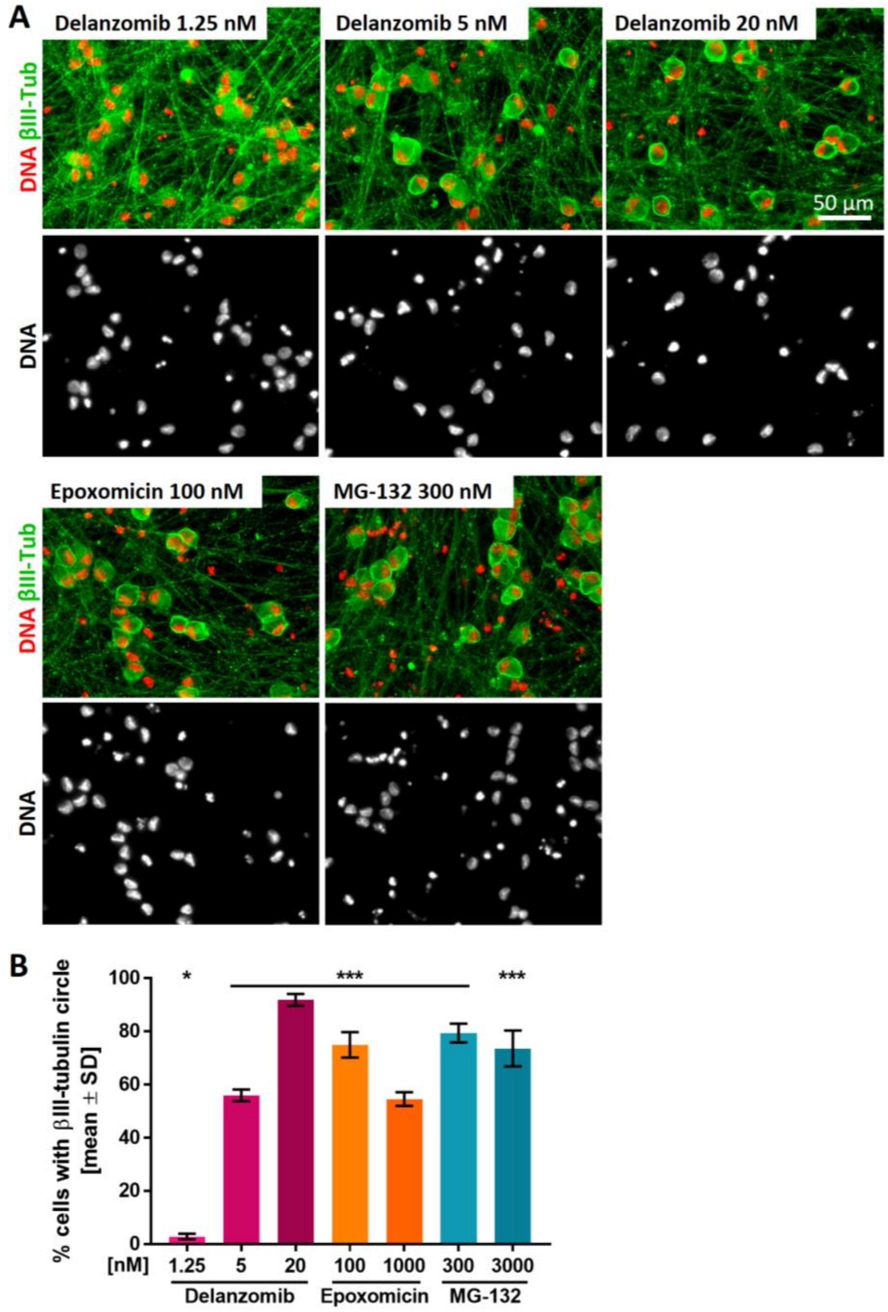


**Fig. S12**
**Holzer et al. 2022**

**Supplementary figure 12: Reorganization of the microtubule structure as a general PI effect**

Sensory neurons (>DoD38) were exposed to the PIs delanzomib, epoxomicin or MG-132 for 24 h before fixation. Representative immunofluorescence images show stainings of nuclei (DNA, red) and βIII-tubulin (βIIITub, green) (see also figure 7G). Composite images are colour coded and single staining images are shown for cell nuclei (DNA). The scale bar is given in the pictures. **(B)** Quantification of cells exhibiting intense, circular βIII-tubulin staining around the cell somata (covering at least 50% of a full circle). Data are given as % of the total cell count (number of viable cell nuclei) and are means ± SD of 2-3 biological replicates. * $p < 0.05$, *** $p < 0.0001$
